# Nanoscale synaptic remodeling at corticostriatal circuits predicts flexible action control

**DOI:** 10.1101/2025.10.30.684628

**Authors:** Vincent Paget-Blanc, Miklós Zöldi, Anna Cavaccini, Vivien Miczán, Barna Dudok, István Katona, Raffaella Tonini

## Abstract

Synaptic plasticity is a fundamental substrate of behavioral adaptation, yet the underlying molecular dynamics remain poorly defined. We tested the hypothesis that, within striatal circuits, flexibility relies on nanoscale remodeling of synaptic machinery coupling anterograde glutamatergic transmission to retrograde endocannabinoid signaling, a process disrupted in states of rigidity and aging. In the dorsolateral striatum, we found cell-type-specific facilitation of metabotropic glutamate receptor 5 (mGlu_5_)-dependent, endocannabinoid-mediated long-term depression at cortico-striatal synapses of indirect pathway neurons in flexible goal-directed behavior, but not after training promoting inflexibility. Stochastic Optical Reconstruction Microscopy (STORM) super-resolution imaging revealed that behavioral adaptation, but not rigidity, is accompanied by increased postsynaptic abundance of mGlu_5_ and diacylglycerol lipase-α (DAGLα), an endocannabinoid-synthesizing enzyme, and presynaptic CB_1_ cannabinoid receptors. In parallel, the nanoscale distance between mGlu_5_ and DAGLα is reduced in postsynaptic spine heads. These nanoscale changes emerged within the time window required for behavioral updating. Intriguingly, the molecular densities of mGlu5, DAGLα, and CB1 receptors predict the strength of behavioral adaptation. In aging mice, these nanoscale changes were absent in association with behavioral rigidity. These findings identify a nanoscale synaptic remodeling mechanism that enables behavioral flexibility and reveal how its failure contributes to rigidity, including that observed in aging.

## Introduction

Automatically executed actions are computationally favorable because they enable faster responses and require reduced cognitive effort. When these actions are represented by learned sequences of acts reinforced through repetition and triggered by cue associations they are defined as habits (1–4). On the other hand, when environmental contingencies change or action outcomes become unpredictable, behavioral adaptation necessitates actions to be under flexible, goal-directed control. Thus, the same inflexible behavioral strategies can be advantageous in stable contexts but disadvantageous in variable ones. Notably, several psychiatric disorders such as obsessive-compulsive disorder and autism spectrum disorder are accompanied by excessive or counter-productive automation of actions (5–8). Moreover, the increased reliance on habits and the related behavioral inflexibility during physiological aging can also become maladaptive if flexibility and adaptation to new circumstances are required. Despite its fundamental physiological and pathological significance, our understanding of the neural processes that underlie the ability or inability to adapt behavioral strategies to changes in action-outcome (A-O) contingency remains limited.

The dorsolateral striatum (DLS) is critically involved in habit formation and expression (9, 10). With instrumental conditioning, over-training in A-O contingency leads to inflexible behavioral control (3, 4, 11) and DLS-specific synaptic modifications of cortico-striatal excitatory synapses (12–17). Moreover, compelling evidence shows that habit learning is accompanied by persistent changes in the strength of glutamatergic neurotransmission (18, 19). On the other hand, the synaptic and molecular mechanisms underlying the inhibition of previously learned behavioral responses to adapt when environmental contingencies change has remained elusive. A potential candidate mechanism involves retrograde endocannabinoid (eCB) signaling at cortico-striatal synapses targeting D_2_ dopamine receptor-expressing indirect pathway (i.e,striatopallidal) medium spiny neurons (iSPNs). Exposure to the psychoactive substance in cannabis, Δ^9^-tetrahydrocannabinol, leads to impaired eCB-mediated signaling in the DLS that is accompanied by habit expression (15). This raises the possibility that molecular mechanisms recruiting retrograde synaptic transmission at cortico-DLS synapses may play a decisive role in adapting behavioral responses to new contingencies.

Retrograde eCB signaling is mediated by the endocannabinoid 2-arachidonoylglycerol (2-AG), a lipid messenger that engages presynaptic CB_1_ cannabinoid receptors (CB_1_Rs) which in turn suppress glutamate release (20, 21). 2-AG is synthesized by the enzyme diacylglycerol lipase-α (DAGL-α) in excitatory synapses (22). At cortico-striatal glutamatergic synapses, 2-AG’s precursor, diacylglycerol, is generated by a biochemical cascade that is primarily elicited by activating group I metabotropic glutamate receptors such as mGlu_5_. The molecular components of this biochemical cascade, as well as mGlu_5_ and DAGL-α, are organized together at the edge of excitatory synapses in the so-called perisynaptic machinery (23). The precise and dynamically regulated nanoscale arrangement of this macromolecular complex is essential for the timely translation of the strength of anterograde neurotransmission into a retrograde feed-back signal. For example, group I mGlu-dependent and eCB-mediated long-term depression is absent at cortico-striatal excitatory synapses in the mouse model of Fragile X syndrome, a monogenic form of autism (24). The average nanoscale distance between mGlu_5_ and DAGL-α is larger than 100 nm at excitatory synapses in these mice that leads to the functional uncoupling between mGlu_5_ and DAGL-α activity. Notably, Fragile X syndrome and other forms of autism are associated with stereotyped behavioral patterns and behavioral inflexibility (25–28). Thus, it is conceivable to hypothesize that synapse-specific nanoscale remodeling of the molecular machinery mediating retrograde eCB signaling is associated with the ability to suppress outdated behavioral responses, thereby enabling changes in action-outcome (A-O) contingency. Moreover, it remains unclear how aging impacts these molecular components at cortico-striatal excitatory synapses in conjunction with age-related behavioral inflexibility.

## Results

We first exploited a behavioral paradigm that measures aspects of behavioral flexibility or inflexibility, respectively, **Fig 1*A***) (15, 29, 30). Mice were subjected to either a short- (*Sh*) or an over-training (*Ov*) session of instrumental conditioning (nose poke for food reward; positive contingency); each training was followed by a reward omission session in which mice were required to inhibit the previously learned behavioral response in the face of a new contingency by refraining from nose poke for food reward (negative contingency). Both groups improved performance throughout the initial training session (active nose poke, ANP/min, session: F_8, 592_ = 167.2, ****p < 0.0001; group: F_3, 74_ = 1.003, p = 0.4; interaction, F_24, 592_ = 1.044, p = 0.4; **Fig 1*B***), and showed similar inactive nose poke (INP) and magazine entry (ME) rates (p ˃ 0.05, **Fig S1*B***). However, only the *Sh*-mice, but not the *Ov*-mice, could adapt to the change in A-O association in the reward omission session (**Fig 1*C***), as indicated by a lower ANP rate computed over the negative contingency (withhold the behavior to maximize reward) compared to the positive contingency: F_1, 31_ = 17.74, ***p = 0.0002, group: F_1, 31_ = 2.436, p = 0.13, contingency X interaction: F_1, 31_ = 1.512, p = 0.23) and by the main group effect of time courses of ANP ratio (time: F_5, 155_ = 10.35, ****p < 0.0001; group: F_1, 31_ = 5.067, *p = 0.0316; interaction: F_5, 155_ = 1.147, p = 0.3382). Moreover, while the two groups earned the same number of pellets during the training session, the *Sh*-mice obtained substantially more pellets than *Ov-*mice during the reward omission session requiring A-O reversal (A-O contingency: F_1, 35_ = 21.82, ****p < 0.0001, group: F_1, 35_ = 14.76, ***p = 0.0005, A-O contingency X interaction: F_1, 35_ = 10.53, **p = 0.0026; **Fig 1*C***). In the four experimental groups, omission sessions yielded similar levels of INP (p ˃ 0.05; **Fig S1*C***). These findings together indicate that the ability to adapt their performance to the change in A-O contingency enabled the *Sh-*mice to accomplish the task more efficiently than the rigid behavioral strategy used by the *Ov-*mice (30).

**Figure 1.**
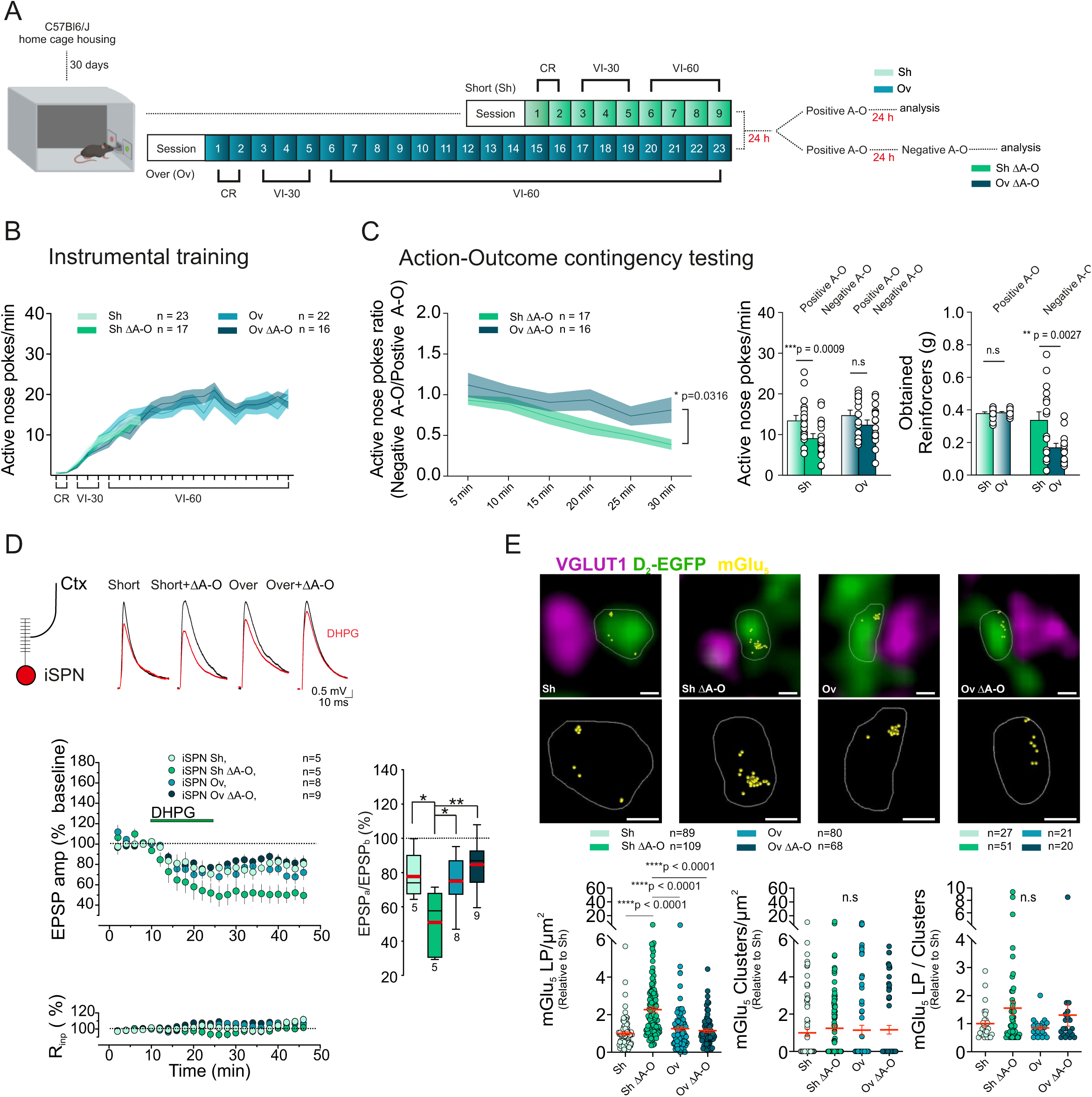
Enhanced mGluR-LTD and increased postsynaptic mGlu5 density during behavioral adaptation. (**A**) Schematic of behavioral paradigm in C57Bl6J mice, CR: continuous reinforcement, VI: variable interval 30 s (VI-30) or 60 s (VI-60). (**B**) C57Bl6J ANP rates during instrumental conditioning in short- and overtrained mice, with or without exposure to a positive and a negative A-O contingency (Sh, n = 23; Ov, n = 22; Sh ΔA-O, n = 17, Ov ΔA-O, n = 18). (**C**) Post-training omission procedure in C57Bl6J. (Left) Time course of ANP ratios (ANP rates under negative A-O/ANP rates under positive A-O) in short- and overtrained mice. (Middle) Comparison of ANP rates between positive and negative A-O contingency in both short- and overtrained mice (Sh n = 17, positive A-O: 13.40 ± 1.29, negative A-O: 9.05 ± 1.12, Sidak ***p = 0.0009; Ov n = 16, positive A-O: 14.76 ± 1.21, negative A-O 12.37 ± 1.17; Sidak p = 0.09). (Right) Number of obtained reinforcers (pellets; g) in the positive- and negative A-O contingency sessions (Sh n = 17, positive A-O: 0.38 ± 0.04, Ov n = 16, positive A-O: 0.34 ± 0.21, Sidak p = 0.9850; Sh negative A-O: 0.39 ± 0.02; Ov negative A-O 0.17 ± 0.10; Sidak **p = 0.0027). (**D**) mGlu-LTD at cortico-iSPN synapses. mGlu-LTD was facilitated in *Sh* ΔA-O-mice compared to all the other training groups (Sh: n = 5 cells, N = 5 mice, 77.8 ± 6.1%; ShΔAO: n = 5 cells, n = 5 mice; Ov: n = 8 cells, N = 5 mice, 50.8 ± 8.7; OvΔAO: n = 8 cells, N = 7 mice, 75.1 ± 5.5; Dunnett’s **p = 0.0068). (**E**) (Top) Combined maximum intensity projections of deconvolved confocal stacks of EGFP-filled iSPN dendrites (green) opposite to VGLUT1-positive axon terminals (magenta) in the DLS of D_2_-EGFP mice, and overlaid mGlu_5_ dSTORM images (yellow); scale bar 200 nm. (Middle) ROIs of the dSTORM images, showing the nanoscale distribution of mGlu_5_ (yellow) in the measured postsynaptic profiles; scale bar 200 nm. Each caption is a representative image from mice subjected to different instrumental training regimes. (Bottom) Quantifications of mGlu_5_ localization points per µm^2^ (Sh, n= 89 ROIs, n = 11 slices, N = 4 mice, 1.00 ± 0.09; Sh ΔA-O, n=109 ROIs, n = 14 slices, N = 4 mice, 2.27 ± 0.13; Ov, n= 80 ROIs, n = 10 slices, N = 4 mice, 1.25 ± 0.12; Ov ΔA-O, n = 68 ROIs, n = 8 slices, N = 4 mice, 1.15 ± 0.10; Dunnett’s, Sh vs. ShΔAO ****p = <0.0001, ShΔAO vs. Ov ****p < 0.0001, ShΔAO vs. OvΔAO ****p < 0.0001); mGlu_5_ cluster numbers per µm^2^ (Sh, n= 89 ROIs, n = 11 slices, n = 4 mice, 1.00 ± 0.20; Sh ΔA-O, n=109 ROIs, n = 14 slices, N = 4 mice, 1.24 ± 0.18; Ov, n= 80 ROIs, n = 10 slices, N = 4 mice, 1.14 ± 0.25; Ov ΔA-O, n = 68 ROIs, n = 8 slices, n = 4 mice, 1.16 ± 0.23); and mGlu_5_ localization points per cluster (Sh, n= 27 ROIs, n = 10 slices, N = 4 mice, 1.00 ± 0.11; Sh ΔA-O, n=51 ROIs, n = 14 slices, N = 4 mice, 1.55 ± 0.25; Ov, n= 21 ROIs, n = 7 slices, N = 4 mice, 0.85 ± 0.07; Ov ΔA-O, n = 20 ROIs, n = 6 slices, N = 4 mice, 1.31 ± 0.39). Data are presented as ratios relative to the averaged values measured in samples from mice subjected to the short training regimen, and are expressed as the mean ± SEM.

The metabotropic glutamate receptor mGlu_5_ has been implicated in memory retention, perseverative behaviors, and extinction learning (31–33). Its activation triggers retrograde eCB signaling that mediates long-term synaptic depression (LTD) at cortico-striatal synapses on the principal striatal projection neurons of the DLS (15, 30, 34, 35). Intriguingly, eCB-mediated LTD is preferentially lost in DLS iSPNs in association with habitual control of behavior induced by exposure to addictive drugs or task overtraining (15, 16). Therefore, we next measured the magnitude of LTD at cortico-iSPN synapses in response to the pharmacological stimulation of mGlu_5_, in acute brain slices obtained from *Sh*-mice and *Ov*-mice before or 30 minutes after the reward omission session (*S*h; *Sh* ΔA-O; *Ov*; *Ov* ΔA-O). The SPNs were identified by their negative resting membrane potential, their firing pattern, and the expression of adenosine receptor A2A (for iSPNs) or Substance P (for dSPNs) as neuronal markers (35). In all four experimental groups, application of the group I mGlu agonist DHPG (100 μM) readily triggered LTD of excitatory postsynaptic potentials (EPSPs) in iSPNs evoked by electrical stimulation of deep cortical layer 5 (**Fig 1*D***). However, mGlu-LTD was substantially facilitated in *Sh* ΔA-O-mice compared to *Sh-*mice, *Ov*-mice and *Ov* ΔA-O-mice (∼50% vs ∼20%). Notably, this effect is cell-type specific, because facilitation of mGluR-LTD was not observed in dSPNs across any of the four groups analyzed (**Fig. S1D**). Thus, updating A-O contingency is accompanied by the facilitation of mGlu-LTD at cortico-iSPN synapses in the DLS.

To elucidate the specific molecular mechanism that accounts for stronger mGlu-LTD in mice that experienced shifting their behavioral strategy, we repeated the behavioral paradigm in four experimental groups of an independent cohort of mice (*Sh*; *Sh* ΔA-O; *Ov*; *Ov* ΔA-O). We took advantage of the Drd2-EGFP BAC transgenic mouse line (D_2_-EGFP) (36) to selectively visualize iSPNs in subsequent molecular analysis. At the behavioral level, these genetically-modified mice recapitulated adaptive responses measured in C57Bl6/J animals (**Fig S1*E-I***). We performed correlated confocal and STORM super-resolution imaging on striatal sections obtained from these mice to measure the nanoscale molecular abundance of mGlu_5_ at cortico-striatal afferent synapses of iSPNs. Immunolabeling for Vesicular Glutamate Transporter 1 (VGLUT1) was used to distinguish cortical glutamatergic afferents from thalamo-striatal and other subcortical excitatory afferents, whereas immunolabeling for EGFP visualized the spine heads of iSPNs receiving VGLUT1-positive glutamatergic excitatory afferents (**Fig 1*E***). The overall number of STORM localization points (LPs) representing the nanoscale position of mGlu_5_ in the EGFP-positive spine heads increased in *Sh* ΔA-O-mice compared to *Sh*-mice, whereas there was no change *Ov*-mice and *Ov* ΔA-O-mice (**Fig 1*E***). In contrast, clustering analysis did not reveal differences among the experimental groups (**Fig 1*E***). These data show that behavioral adaptation to a new task demand and stronger mGlu-LTD occur in parallel to an overall increase in the density of postsynaptic mGlu_5_-immunolabeling opposite of cortico-striatal afferents of iSPNs, but there is no major change in the nanoscale distribution of mGlu_5_ receptors.

Because mGlu_5_ receptor activity is functionally coupled with the eCB-producing enzyme diacylglycerol lipase-α (DAGLα) in the postsynaptic spine head, we tested whether flexible behavioral performance and stronger mGlu-LTD are accompanied by an upregulation of the molecular machinery for eCB production and/or the number of presynaptic CB_1_ cannabinoid receptors (CB_1_Rs) on cortico-striatal axon terminals targeting iSPNs. Correlated confocal and STORM microscopy revealed an increased number of localization points representing DAGLα in spine heads of iSPNs receiving VGLUT1-positive afferents exclusively in *Sh* ΔA-O-mice in comparison with *Sh*-mice, *Ov*-mice and *Ov* ΔA-O-mice (**Fig 2*A* and *B***). Furthermore, more DAGLα clusters were observed in spine heads in *Sh* ΔA-O-mice and these clusters consisted of almost twice as many individual DAGLα-representing localization points as those in the other groups (**Fig 2*B***).

**Figure 2.**
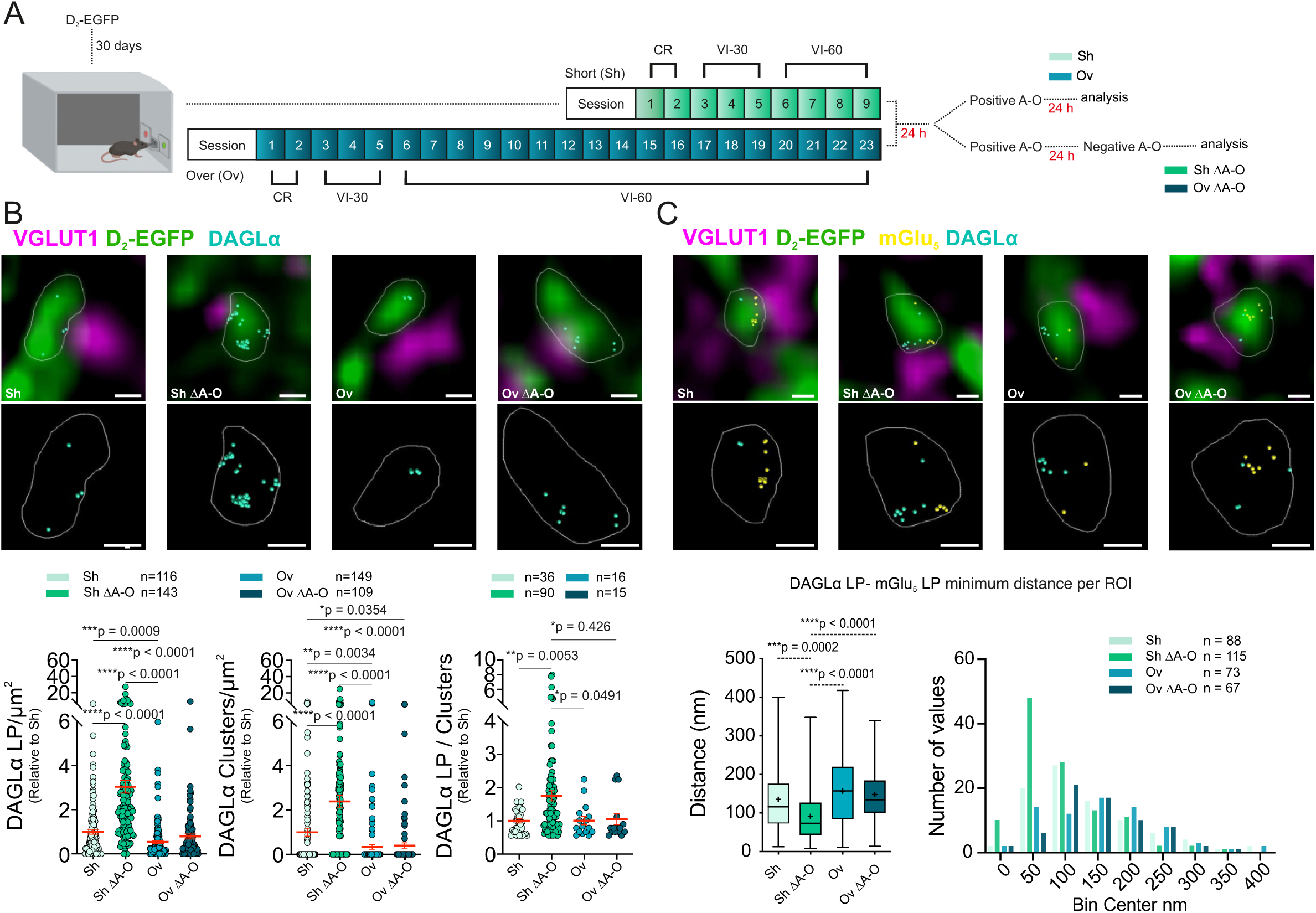
Flexible behavioral performance is accompanied by nanoscale reorganization of postsynaptic mGluR_5_ and DGLα. (**A**) Schematic of the behavioral paradigm in D_2_–EGFP mice, CR: continuous reinforcement, VI: variable interval 30 s (VI-30) or 60 s (VI-60). (**B**) (Top) Combined maximum intensity projections of deconvolved confocal stacks of EGFP-filled iSPN dendrites opposed to VGLUT1 boutons in the DLS of D_2_-EGFP mice, and overlaid DAGLα dSTORM images (cyan); scale bar 200 nm. (Middle) ROIs of the dSTORM images showing the nanoscale distribution of DAGLα in the postsynaptic profiles receiving the VGLUT1-positive cortico-striatal afferent synapses. Each caption is a representative image from mice subjected to different instrumental training regimes; scale bar 200 nm. (Bottom) Quantification of localization points representing DAGLα per µm^2^ (Sh, n = 116 ROIs, n = 8 slices, N = 3 mice, 1 ± 0.11; Sh ΔA-O, n = 143 ROIs, n = 10 slices, n = 4 mice, 3.03 ± 0.28; Ov, n = 149 ROIs, n = 11 slices, N = 4 mice, 0.54 ± 0.07; Ov ΔA-O, n = 109 ROIs, n = 9 slices, N = 3 mice, 0.78 ± 0.11; Dunnett’s, Sh vs. Sh ΔA-O ****p < 0.0001, Sh vs. Ov ***p = 0.0009 Sh ΔA-O vs. Ov ****p < 0.0001, ShΔAO vs. OvΔAO ****p < 0.0001); DAGLα cluster number per µm^2^ (Sh, n = 116 ROIs, n = 8 slices, N = 3 mice, 1 ± 0.18; Sh ΔA-O, n = 143 ROIs, n = 10 slices, N = 4 mice, 2.39 ± 0.27; Ov, n = 149 ROIs, n = 11 slices, N = 4 mice, 0.34 ± 0.1; Ov ΔA-O, n = 109 ROIs, n = 9 slices, N = 3 mice, 0.4 ± 0.12; Dunnett’s, Sh vs. Sh ΔA-O ****p < 0.0001, Sh vs. Ov ** p = 0.0034, Sh vs. Ov ΔAO *p = 0.0354, Sh ΔA-O vs. Ov ****p < 0.0001, ShΔAO vs. OvΔAO ****p < 0.0001); and DAGLα localization points per cluster (Sh, n = 36 ROIs, n = 7 slices, N = 3 mice, 1 ± 0.06; Sh ΔA-O, n = 90 ROIs, n = 10 slices, N = 4 mice, 1.76 ± 0.15; Ov, n = 16 ROIs, n = 9 slices, N = 4 mice, 1.01 ± 0.12; Ov ΔA-O, n = 15 ROIs, n = 6 slices, N = 3 mice, 1.06 ± 0.17; Dunnett’s, Sh vs. Sh ΔA-O **p = 0.0053, Sh ΔA-O vs. Ov *p = 0.0491, ShΔAO vs. OvΔAO *p = 0.0426). Data are presented as ratios relative to the averaged values measured in samples from mice subjected to the short training regimen, and are expressed as the mean ± SEM. (**C**) (Top) Combined maximum intensity projections of deconvolved confocal stacks of EGFP-filled iSPN dendrites (green) from D_2_-EGFP mice opposite to VGLUT1-positive corticostriatal axon terminals (magenta), and overlaid mGlu_5_ and DAGLα dSTORM images (yellow, cyan; respectively); scale bar 200 nm. (Middle) ROIs of the dSTORM images depicting the nanoscale distribution of both mGlu_5_ (yellow) and DAGLα (cyan) in the postsynaptic domain. Each caption is a representative image from mice subjected to different instrumental training regimes; scale bar 200 nm. (Bottom left) Quantification of the nearest neighbor distance between DAGLα-representing localization points and mGlu_5_-representing localization points (Sh, n = 88 distances, N= 4 mice, 135.80 ± 8.80; Sh ΔA-O, n = 115 distances, N = 4 mice, 91.43 ± 6.12; Ov, n = 73 distances, N = 4 mice, 156.90 ± 10.24; Ov ΔA-O, n = 67 distances, N = 4 mice, 147.90 ± 8.78; Dunnett’s, Sh vs. ShΔAO ***p = 0.0002, ShΔAO vs. Ov ****p = <0.0001, ShΔAO vs. OvΔAO ****p < 0.0001). (Bottom right) Distribution of the nearest neighbor distance between DAGLα localization points to mGlu_5_ localization points in mice subjected to different instrumental training regimes. Distance distribution counts are binned per 50 nm. All data are expressed as mean ± SEM except for binned data (expressed as number of values).

Both mGlu_5_ and DAGLα are transiently recruited to the perisynaptic machinery via binding to the anchoring protein Homer, and their nanoscale vicinity is required for their functional coupling (24). By exploiting the diffraction-unlimited localization precision of dual-channel STORM super-resolution imaging, we measured the nearest neighbor distances between mGlu5 and DAGLα. In Sh-mice, Ov-mice, and Ov ΔA-O-mice, these distances averaged between 127 and 167 nm. In contrast, in Sh ΔA-O-mice, the nearest neighbor distances were substantially reduced, with a mean at 91 nm (**Fig 2*C***). These findings together demonstrate that the ability to engage specific nanoscale molecular changes in the spine heads of iSPNs accompaines flexible behavioral responses, a capacity that is absent in overtrained, inflexible mice.

Because the upregulation of the perisynaptic molecular machinery indicates that the translation of anterograde glutamatergic transmission to retrograde eCB signaling may be more effective at cortico-striatal afferents of iSPNs when a behavioral strategy is updated, we next asked whether the molecular density of presynaptic CB_1_ receptors mirrors the postsynaptic molecular adaptations. Correlated confocal and STORM super-resolution imaging of cortico-striatal boutons terminating on the spine heads of iSPNs uncovered that the number of localization points representing presynaptic CB_1_ receptors is almost 4 times larger in *Sh* ΔA-O-mice than in *Sh*-mice, *Ov*-mice and *Ov* ΔA-O-mice (**Fig 3*A*-*B***). In addition, the density of CB_1_ receptor clusters was also substantially higher only in *Sh* ΔA-O-mice while the number of individual localization points per cluster did not vary (**Fig 3*B***). Taken together the STORM imaging data reveal a consistent increase in the molecular abundance of the presynaptic and postsynaptic molecular players of retrograde eCB signaling at cortico-striatal afferent of iSPNs in conjunction with stronger mGlu-LTD and the ability to update changes in A-O contingencies. While there was a clear segregation in the ability or inability of *Sh* ΔA-O-mice and *Ov* ΔA-O-mice, respectively, to adapt to the new task demands (**Fig 4*A***), both experimental groups exhibited substantial individual variation in performance. By exploiting the broad range of active nose pokes during the reward omission session (**Fig 1*C***), we posed the question of whether the nanoscale molecular adaptation at cortico-striatal afferents to iSPNs predicts individual differences in behavioral flexibility. We calculated the inverse ratio (ANP negative A-O contingency/ANP positive A-O contingency) as the adaptive index for each animal. Notably, the number of postsynaptic mGlu_5_-positive localization points strongly and positively correlated with the adaptive index. In contrast, neither the cluster number nor the number of localization points in clusters scaled with the adaptive index (**Fig 4*B***). Moreover, the adaptive index of an individual animal was also reliably predicted by the overall number and the cluster number of DAGLα located postsynaptically at cortico-striatal synapses of iSPNs (**Fig 4*C***). Similarly, most molecular parameters of CB_1_-immunolabeling exhibited substantial positive correlation with the adaptive index of individual animals (**Fig 4*D***). These findings suggest that the density of the key molecular elements of retrograde eCB signaling at cortico-striatal inputs of iSPNs co-vary with the ability of an animal to adapt to the new contingency.

**Figure 3.**
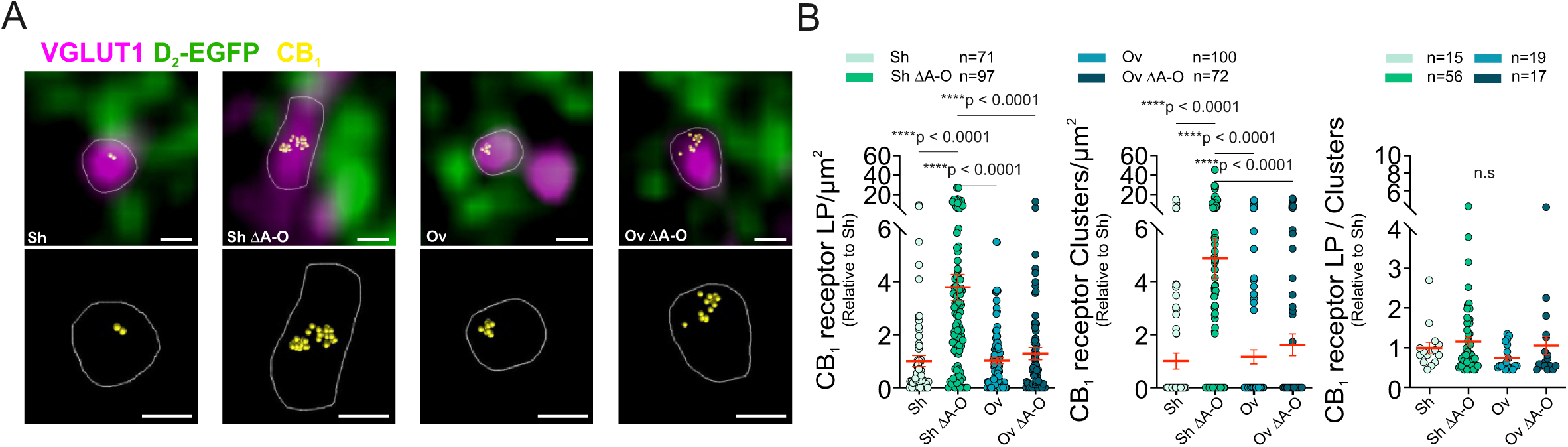
In flexible mice, the molecular density of presynaptic CB_1_ receptors mirrors the postsynaptic molecular adaptations. (**A**) (Top) Combined maximum intensity projections of deconvolved confocal stacks of EGFP-filled iSPN dendrites (green) that are located opposite to VGLUT1-positive boutons (magenta) and overlaied dSTORM images of the nanoscale distribution of CB_1_ receptors (yellow) in the axon terminals; scale bar 200 nm. (Bottom) ROI of dSTORM images; scale bar 200 nm. Each caption is a representative image in mice subjected to different instrumental training regimes. (**B**) Quantification of CB_1_ receptor localization points per µm^2^ (Sh, n = 71 ROIs, n = 6 slices, N = 3 mice, 1 ± 0.21; Sh ΔA-O, n = 97 ROIs, n = 8 slices, N = 4 mice, 3.79 ± 0.49; Ov, n = 100 ROIs, n = 8 slices, N = 4 mice, 1.02 ± 0.11; Ov ΔA-O, n = 72 ROIs, n = 6 slices, N = 3 mice, 1.28 ± 0.23; Dunnett’s, Sh vs. Sh ΔAO ****p < 0.0001, Sh ΔAO vs. Ov ****p < 0.0001, Sh ΔAO vs. Ov ΔAO ****p < 0.0001); CB_1_ receptor cluster number per µm^2^ (Sh, n = 71 ROIs, n = 6 slices, N = 3 mice, 1 ± 0.3; Sh ΔA-O, n = 97 ROIs, n = 8 slices, N = 4 mice, 4.87 ± 0.74; Ov, n = 100 ROIs, n = 8 slices, N = 4 mice, 1.16 ± 0.27; Ov ΔA-O, n = 72 ROIs, n = 6 slices, N = 3 mice, 1.61 ± 0.42; Dunnett’s, Sh vs. Sh ΔAO ****p < 0.0001, Sh ΔAO vs. Ov ****p < 0.0001, Sh ΔAO vs. Ov ΔAO ****p < 0.0001); CB_1_ receptor localization points per cluster (Sh, n = 15 ROIs, n = 4 slices, N = 3 mice, 1 ± 0.14; Sh ΔA-O, n = 56 ROIs, n = 8 slices, N = 4 mice, 1.16 ± 0.11; Ov, n = 19 ROIs, n = 6 slices, N = 4 mice, 0.73 ± 0.07; Ov ΔA-O, n = 17 ROIs, n = 6 slices, N = 3 mice, 1.06 ± 0.23). Data are presented as ratios relative to the averaged values measured in samples from mice subjected to the short training regimen, and are expressed as mean ± SEM.

**Figure 4.**
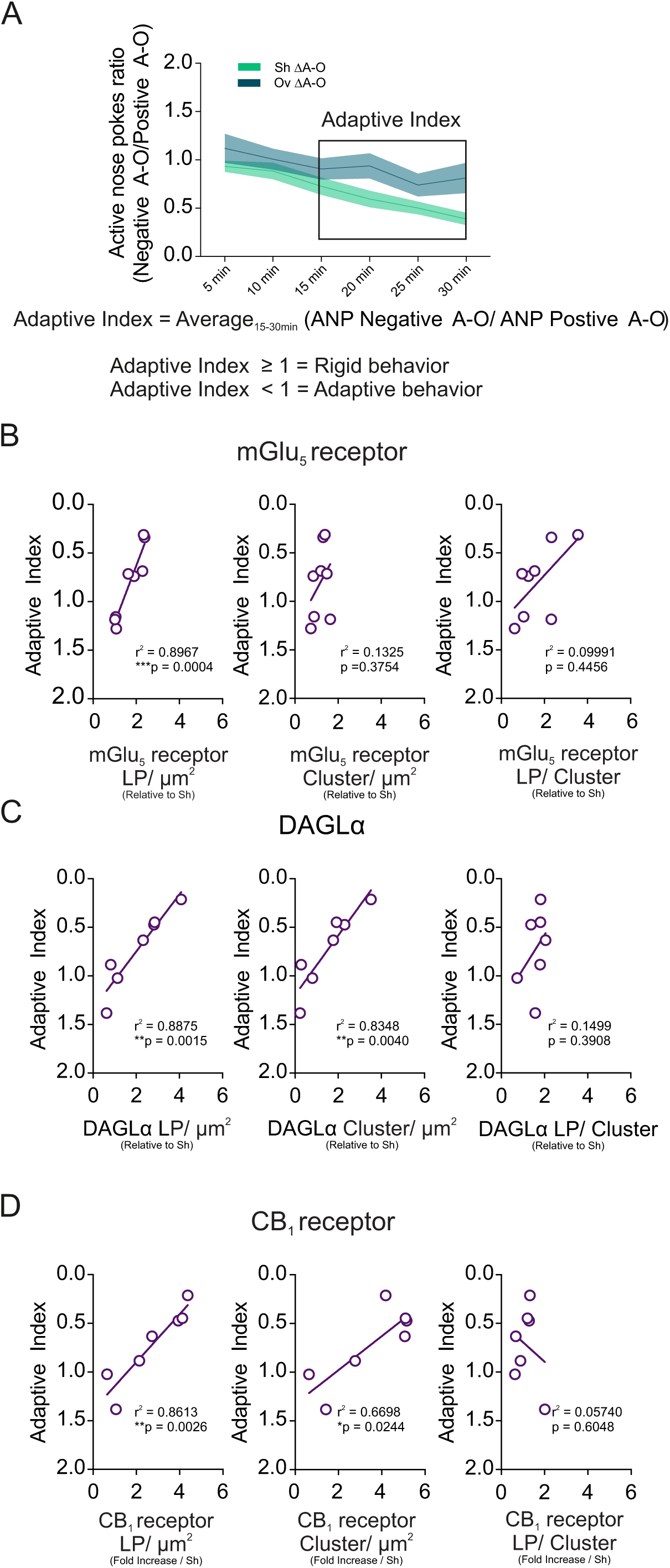
Nanoscale molecular adaptation at cortico-striatal afferents to iSPNs predicts adaptive behavioral performance. (**A**) Schematic presentation of the definition of the Adaptive Index behavioral parameter. (**B**) Correlation between the Adaptive Index and mGlu_5_-representing localization points (left), mGlu_5_-containing clusters (middle), mGlu_5_ localization points per clusters (right). All localization points data is shown relative to the data obtained Sh training regime group (two-tailed Pearson’s correlation coefficient: LP, ***p = 0.0004, r^2^ = 0.8967; Clusters, p = 0.3754, r^2^ = 0.1325; LP / Clusters, p = 0.4456, r^2^ = 0.09991). Each dot represents values from an individual mice. (**C**) Correlation between the Adaptive Index and DAGLα-representing localization points (left), DAGLα-containing clusters (middle), DAGLα localization points per clusters (right). All localization points data is shown relative to the data obtained Sh training regime group (two-tailed Pearson’s correlation coefficient: LP, **p = 0.0015, r^2^ = 0.8875; Clusters, **p = 0.0040, r^2^ = 0.8348; LP / Clusters, p = 0.3908, r^2^ = 0.1499). Each dot represents values from an individual mice. (**D**) Correlation between the Adaptive Index and CB_1_ receptor-representing localization points (left), CB_1_ receptor-containing clusters (middle), CB_1_ receptor localization points per clusters (right). All localization points data is shown relative to the data obtained Sh training regime group (two-tailed Pearson’s correlation coefficient: LP, **p = 0.0026, r^2^ = 0.8613; Clusters, *p = 0.0244, r^2^ = 0.6698; LP / Clusters, p = 0.6048, r^2^ = 0.05740). Each dot represents values from an individual mice.All data is expressed as mean ± SEM.

Diminished ability to update a previously acquired behavior is a characteristic feature of aging. We reasoned that if the increased number of the postsynaptic DAGLα enzymes and the presynaptic CB_1_ receptors represent a molecular signature for the ability to change behavioral strategy then the molecular adaptations observed in young animals in association with an altered A-O contingency may be compromised in old mice. Mice were housed for one year and then underwent the same short-training session as young mice housed for 30 days (**Fig 5*A***). The two age groups performed the same number of ANPs during the training session indicating similar ability of instrumental learning (**Fig 5*B***). Training and reward omission sessions in the different experimental groups yielded similar levels of INPs and ME (p ˃ 0.05; **Fig S2*B-C***). On the other hand, old animals exhibited a large variability in their ability to refrain from nose poke for food reward when subjected to omission (**Fig 5*C***). This variability matched the range of short-trained and over-trained animals. The large variance in the adaptive index of aged mice made their assignement into two groups possible. Using k-nearest neighbors clustering analysis (kNN), we categorized the behavioral performance from *Sh* ΔA-O- and *Ov* ΔA-O-mice into adaptive and rigid. Those aged mice *(Sh* aged) that had an adaptive index and number of obtained rewards in the range of *Sh* ΔA-O young animals were assigned to the group of *Sh* aged ΔA-O Adaptive-mice, whereas those resembling *Ov* ΔA-O mice were classified as *Sh* aged ΔA-O Rigid-mice (**Fig 5*C*** and ***D***). While the former group could readily refrain from nose poking for food reward upon omission, the latter group did not show a change in behavioral strategy (**Fig 5*D***). Notably, under basal conditions, the molecular parameters of DAGLα-immunolabeling were similar between young and aged short-trained animals (**Fig 5*E***). On the other hand, correlated confocal and STORM microscopy on cortico-striatal excitatory synapses on the spine heads of iSPNs revealed that the overall number of localization points representing DAGLα, and the number of DAGLα-containing clusters, were both strongly increased in *Sh* aged ΔA-O Adaptive-mice compared to all other groups, whereas *Sh* aged ΔA-O Rigid-mice did have a reduced DAGLα-immunolabeling following ΔA-O (**Fig 5*E***). Similarly, localization points representing presynaptic CB_1_Rs in cortico-striatal axon terminals exhibited a robust increase in *Sh* aged ΔA-O Adaptive-mice, and was reduced in *Sh* aged ΔA-O Rigid-mice compared to young short-trained animals (**Fig. 5*F***). Taken together, molecular adaptation of the retrograde eCB signaling pathway accompanies the ability to update changes in action-outcome (A-O) contingencies even in old mice, whereas the same molecular plasticity phenomenon is absent in old mice exhibiting behavioral inflexibility.

**Figure 5.**
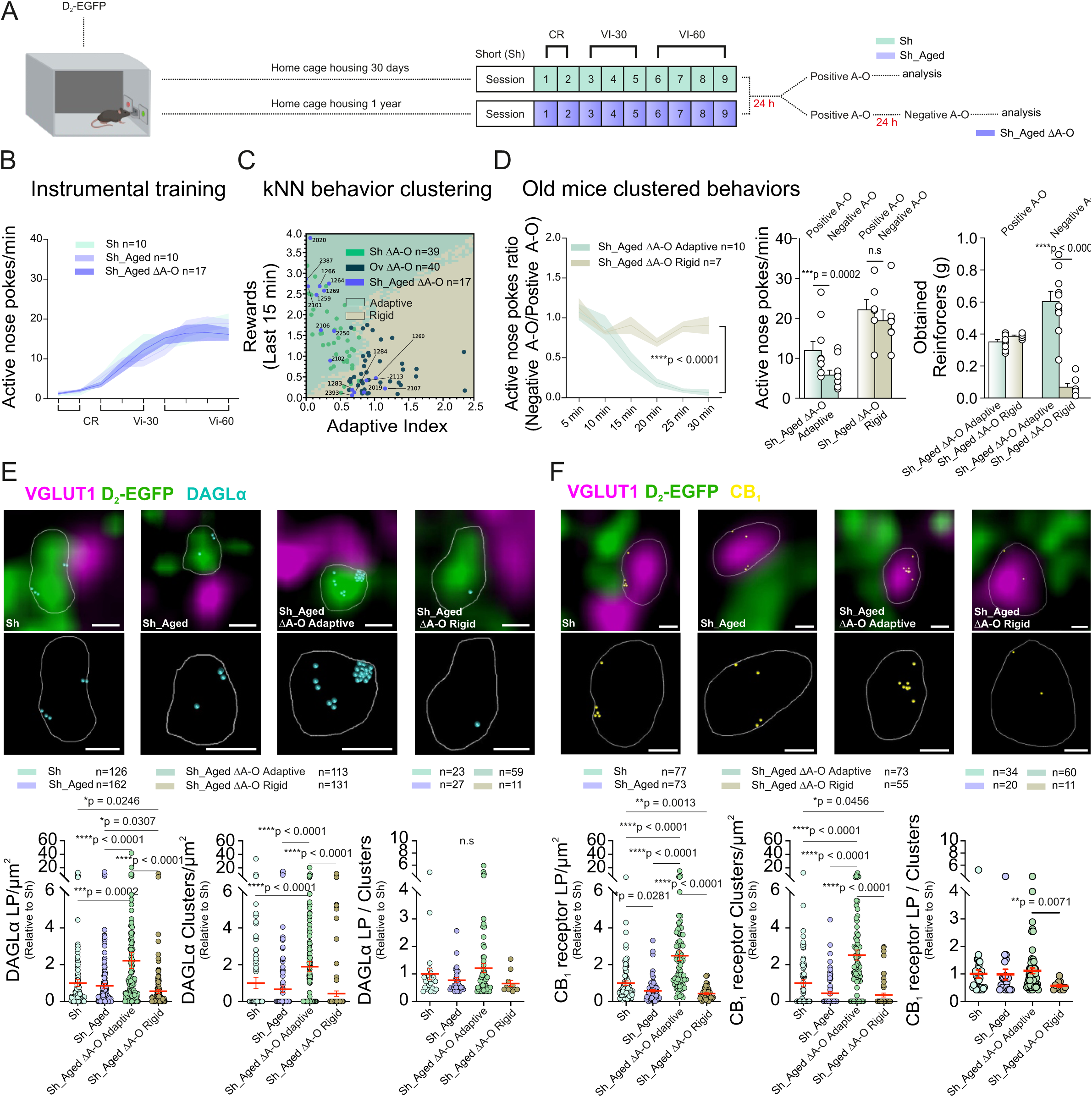
mGluR_5_–DGL nanoscale plasticity is impaired in aged mice with inflexible behavior. (**A**) Schematic summary of the behavioral paradigm, CR: continuous reinforcement, VI: variable interval 30 s (VI-30) or 60 s (VI-60). (**B**) ANP rates during instrumental conditioning in the four mice cohorts (Sh, n = 10; Sh_Aged, n = 10; Sh_Aged ΔA-O, n = 17). (**C**) k-Nearest Neighbors behavior clustering of aged mice subjected to a short training followed by ΔA-O (Sh_Aged ΔA-O, n = 17). The matrice was created by aggregating data from the last 15 minutes of ANP ratios (ANP rates under negative A-O/ANP rates under positive A-O) and reward ratios (reward rates under negative A-O/rewards rates under positive A-O) computed in short-trained adaptive and over-trained rigid mice using both the subjects from this study and additional data available in the laboratory (Sh ΔA-O, n = 38; Ov ΔA-O, n = 40). The k factor selected for clustering was eight, and the step size for the matrice was 0.05. (**D**) Post-training omission procedure. (Left) Time course of ANP ratios (ANP rates under negative A-O/ANP rates under positive A-O) of short-trained mice housed in home cages for one year clustered through kNN (Sh_Aged ΔA-O Flexible, n = 10; Sh_Aged ΔA-O Inflexible, n = 7). (Middle) Comparison of ANP rates between positive and negative A-O contingency in both short-trained aged mice (Sh_Aged Flexible n = 10, positive A-O: 12.05 ± 2.13, negative A-O: 5.89 ± 1.13, Sidak ***p = 0.0002; Sh_Aged inflexible n = 7, positive A-O: 22.20 ± 2.46, negative A-O 19.48 ± 2.60; Sidak p = 0.1437). (Right) Number of obtained reinforcers (pellets; g) in the positive-and negative A-O contingency sessions (Sh_Aged ΔAO Flexible n = 10, positive A-O: 0.35 ± 0.01, Sh_Aged ΔAO Inflexible n = 7, positive A-O: 0.39 ± 0.01, Sidak p = 0.772; Sh_Aged ΔAO Flexible negative A-O: 0.60 ± 0.06; Sh_Aged ΔAO Inflexible negative A-O 0.07 ± 0.02; Sidak ****p < 0.0001). (**E**) (Top) Combined maximum intensity projections of deconvolved confocal stacks of EGFP-filled iSPN dendrites (green) opposite to VGLUT1-positive boutons (magenta) and overlayed DAGLα dSTORM image (cyan); scale bar 200 nm. (Middle) ROI of dSTORM images showing the nanoscale distribution of DAGLα (cyan) in the postsynaptic domain of the same synapse; scale bar 200 nm. Each caption represents young adult or aged mice subjected to short instrumental training regime and expressing either behavioral flexibility or inflexibility. (Bottom left) Quantification of DAGLα localization points per µm^2^ relative to the short training regime condition from young adult mice (Sh, n = 126 ROIs, n = 11 slices, n = 4 mice, 1 ± 0.18; Sh_Aged, n = 162 ROIs, n = 10 slices, n = 4 mice, 0.85 ± 0.13; Sh_Aged ΔAO Flexible, n = 113 ROIs, n = 9 slices, N = 3 mice, 2.22 ± 0.44; Sh_Aged ΔAO Inflexible, n = 131 ROIs, n = 9 slices, n = 3 mice, 0.56 ± 0.11; Dunnett’s, Sh vs. Sh_Aged ΔA-O Flexible ***p = 0.0002, Sh vs. Sh_Aged ΔA-O Inflexible *p = 0.0246, Sh_Aged vs. Sh_Aged ΔA-O Flexible ****p < 0.0001; Sh_Aged vs. Sh_Aged ΔA-O Inflexible *p = 0.0307, Sh_Aged ΔA-O Flexible vs. Sh_Aged ΔA-O inflexible ****p < 0.0001). (Bottom middle) Quantification of DAGLα cluster number per µm^2^ relative to the young adult short training regime condition (Sh, n = 126 ROIs, n = 11 slices, n = 4 mice, 1 ± 0.31; Sh_Aged, n = 162 ROIs, n = 10 slices, n = 4 mice, 0.66 ± 0.18; Sh_Aged ΔAO Flexible, n = 113 ROIs, n = 9 slices, N = 3 mice, 1.89 ± 0.28; Sh_Aged ΔAO Inflexible, n = 131 ROIs, n = 9 slices, n = 3 mice, 0.43 ± 0.15; Dunnett’s, Sh vs. Sh_Aged ΔA-O Flexible ****p < 0.0001, Sh_Aged vs. Sh_Aged ΔA-O Flexible ****p < 0.0001, Sh_Aged ΔA-O Flexible vs. Sh_Aged ΔA-O inflexible ****p < 0.0001). (Bottom right) Quantification of DAGLα localization points per cluster relative to the young adult short training regime condition (Sh, n = 23 ROIs, n = 10 slices, n = 4 mice, 1 ± 0.21; Sh_Aged, n = 27 ROIs, n = 10 slices, n = 4 mice, 0.77 ± 0.09; Sh_Aged ΔAO Flexible, n = 59 ROIs, n = 9 slices, N = 3 mice, 1.21 ± 0.16; Sh_Aged ΔAO Inflexible, n = 11 ROIs, n = 7 slices, n = 3 mice, 0.65 ± 0.11). Data are presented as ratios relative to the averaged values measured in samples from mice subjected to the short training regimen, and are expressed as the mean ± SEM. (**F**) (Top) Combined maximum intensity projections of deconvolved confocal stacks of EGFP-filled iSPN dendrites from D_2_-EGFP mice opposed to VGLUT1 boutons and overlayed CB_1_ receptor dSTORM images (yellow); scale bar 200 nm. (Midle) dSTORM images showing the nanoscale distribution of CB_1_ receptor (yellow) in ROIs of the same synapse. Each caption represents young adult or aged mice subjected to a short instrumental training regime and expressing behavioral flexibility or inflexibility; scale bar 200 nm. (Bottom left) Quantification of CB_1_ receptor localization points per µm^2^ expressed relative to the short training regime condition from young adult mice (Sh, n = 77 ROIs, n = 8 slices, N = 4 mice, 1.00 ± 0.12; Sh_Aged, n = 73 ROIs, n = 8 slices, N = 4 mice, 0.58 ± 0.07; Sh_Aged ΔAO Flexible,n = 73 ROIs, n = 9 slices, N = 3 mice, 2.49 ± 0.26; Sh_Aged ΔAO Inflexible, n = 55 ROIs, n = 9 slices, N = 3 mice, 0.43 ± 0.05; Dunnett’s, Sh vs. Sh_Aged, *p = 0.0281 Sh vs. Sh_Aged ΔA-O Flexible ****p < 0.0001, Sh vs. Sh_Aged ΔA-O Inflexible **p = 0.0013, Sh_Aged vs. Sh_Aged ΔA-O Flexible ****p < 0.0001, Sh_Aged ΔA-O Flexible vs. Sh_Aged ΔA-O inflexible ****p < 0.0001). (Bottom middle) Quantification of CB_1_R cluster number per µm^2^ expressed relative to the young adult short training regime condition (Sh, n = 77 ROIs, n = 8 slices, N = 4 mice, 1.00 ± 0.21; Sh_Aged, n = 73 ROIs, n = 8 slices, N = 4 mice, 0.44 ± 0.10; Sh_Aged ΔAO Flexible, n = 73 ROIs, n = 9 slices, N = 3 mice, 2.52 ± 0.28; Sh_Aged ΔAO Inflexible, n = 55 ROIs, n = 9 slices, N = 3 mice, 0.34 ± 0.11; Dunnett’s, Sh vs. Sh_Aged ΔA-O Flexible ****p < 0.0001, Sh vs. Sh_Aged ΔA-O Inflexible *p = 0.0456, Sh_Aged vs. Sh_Aged ΔA-O Flexible ****p < 0.0001, Sh_Aged ΔA-O Flexible vs. Sh_Aged ΔA-O inflexible ****p < 0.0001). (Bottom right) Quantification of CB_1_ receptor localization points per cluster number expressed relative to the young adult short training regime condition (Sh, n = 34 ROIs, n = 8 slices, N = 4 mice, 1 ± 0.14; Sh_Aged, n = 20 ROIs, n = 8 slices, N = 4 mice, 0.98 ± 0.19; Sh_Aged ΔAO Flexible,n = 60 ROIs, n = 9 slices, N = 3 mice, 1.11 ± 0.1; Sh_Aged ΔAO Inflexible, n = 11 ROIs, n = 6 slices, N = 3 mice, 0.57 ± 0.05; Dunnett’s, Sh_Aged ΔA-O Flexible vs. Sh_Aged ΔA-O inflexible **p = 0.0071. Data are presented as ratios relative to the averaged values measured in samples from mice subjected to the short training regimen, and are expressed as the mean ± SEM.

## Discussion

Retrograde eCB signaling is critical for various form of synaptic plasticity throughout the brain, yet the specific role of its strength in supporting circuit computation during behavior remains unclear. We show that flexible behavioral control is accompanied by enhanced eCB-mediated LTD at corticostriatal synapses and it is positively correlated with the nanoscale remodeling of the underlying molecular components of retrograde eCB signaling. In contrast, behavioral inflexibility, evident in overtrained mice, is characterized by the lack of these nanoscale adaptations. Similarly, aging mice exhibiting behavioral rigidity fail to show nanoscale plasticity in postsynaptic DAGLα, and presynaptic CB_1_ levels.

Dynamic modulation of the strength of anterograde neurotransmission has fundamental importance in brain circuit computations. For example, consolidation of skill learning is attributed to long-lasting potentiation of cortical glutamatergic transmission in the DLS (18, 37–39). Moreover, the underlying synaptic adaptations are pathway-specific, occur predominantly in D2-expressing iSPNs and have a postsynaptic (but not presynaptic) site of expression for plasticity (18). In contrast, the mechanisms, and the behavioral significance of dynamic changes in the strength of retrograde synaptic signaling have remained much less understood. An ideal paradigm to study these questions in striatal circuits is group I mGlu-dependent, eCB-mediated long-term depression in which the origin of plasticity is postsynaptic, whereas the site of expression is presynaptic (40, 41). mGlu-dependent, eCB-mediated long-term depression in the striatum is known to be associated with a number of physiological and pathological conditions (24, 42–45). Accordingly, we found that the capability of eCBs to depress synaptic transmission in response to mGlu1/5 activation is substantially facilitated at cortico-striatal synapses of DLS iSPNs, but not of dSPNs, in association with the ability of the animals to update actions in response to a shift in action-outcome contingency. In parallel, our data revealed a significant increase in the synaptic abundance of the three key molecular players (mGlu_5_, DGLα and CB_1_ receptors) of mGlu-dependent, eCB-mediated LTD in behaviorally flexible mice. These consistent molecular changes are likely account for the facilitated LTD, raising the possibility of optimized molecular coupling capable of supporting adaptive suppression of outdated actions; notably, this mechanism appears compromised in both ovetrained and aging mice exhibiting behavioral inflexibility.

Our combined approach using correlated confocal microscopy and STORM super-resolution imaging provided critical insights into this synaptic remodeling. Confocal microscopy enables the cell-type-specific delineation of specific subcellular compartments, whereas STORM super-resolution imaging is a quantitative approach to measure molecular distributions within the defined subcellular contexts at the nanoscale level (46, 47). It is important to obtain molecular information below the diffraction limit of traditional optical microscopes, because the nanodomain-specific position of specific signaling molecules define their specific synaptic function and malfunction (48–52). For example, mGlu_5_-dependent, eCB-mediated LTD is absent at corticostriatal synapses in a mouse model of Fragile X syndrome, a monogenic form of autism. The functional uncoupling between anterograde glutamatergic signaling and retrograde endocannabinoid production is due to the nanoscale level increase in the distance between mGlu_5_ receptors and the endocannabinoid 2-AG-synthesizing enzyme DGLα (24). In contrast, we found that the nanoscale decrease in the distance among these molecular players also strongly corelated with the behavioral adaptation capacity of individual animals. Notably, this nanoscale remodeling takes place within the time window required for a mouse to update action-outcome contingencies (∼ 30 min). This indicates that these molecular processes are not only rapid but temporally aligned with the neuronal computations required for behavioral adapatation. As a consideration, molecular distributions in the perfused tissue only represent a snapshot of otherwise continuously moving proteins in the plasma membrane.

In accordance, both mGlu_5_ receptors and CB_1_ receptors have homogeneous distribution along the postsynaptic and presynaptic surface, respectively, that reflect their dynamic motion (52–54). A portion of these signaling proteins becomes confined transiently to their functionally relevant nanodomain (52, 55). Thus, our anatomical findings of the increased abundance of mGlu_5_, DGLα and CB_1_ receptors in the postsynaptic and presynaptic compartments indicate that the probability of their functional coupling is facilitated in association with flexible control of behavior. The specific molecular mechanisms of this coordinated postsynaptic and presynaptic molecular plasticity remain unclear. Trans-synaptic adhesion molecules such as the neuroligin-neurexin complex represent potential candidates for this regulation. For example, neurexin1α loss in the striatum has been shown to enhance eCB-mediated depression at cortico-striatal iSPN synapses (56). While the regulatory mechanisms require further studies, our findings demonstrate for the first time that the molecular plasticity mechanisms controlling the strength of retrograde eCB signaling in the DLS show conceptual similarity (such as nanoscale level changes in receptor numbers) to the synaptic plasticity rules governing the strength of anterograde neurotransmission (57, 58).

The DLS is traditionally associated with the gradual encoding of stimulus-response associations through the repetition of behavior, ultimately leading to habitual actions (8, 10, 11, 59–65). Our omission procedure explicitly evaluates behavioral flexibility in short-trained mice and inflexibility in overtrained mice by testing their ability or inability to suppress previously reinforced actions upon contingencies shift (15, 29, 30, 66, 67). This task critically engages glutamatergic signaling in the DLS (15, 30), including mGluR5-mediated intracellular pathways (17). Unlike tasks associated with associative fronto-DMS circuits that require learning novel actions (68, 69), the omission procedure requires animals to withhold a behavior to maximize reward (70). Consistent with updated models of motor encoding, dSPNs are congruently activated to support ongoing behavior. In contrast, the majority of iSPNs tuned to ongoing behavior remain silent, whereas those to alternative or competing behaviors become active to suppress suboptimal actions (13, 71). A plausible scenario is that adaptive suppression in short-trained, flexible mice might depend on the transient silencing of iSPNs tuned to the now-outdated behavior. Enhanced eCB-mediated depression of corticostriatal synapses onto this iSPN ensemble may enable selective suppression, thereby facilitating disengagement from the previous response and supporting behavioral switching. Compared to other inhibitory neuromodulators such as adenosine and nitric oxide, the retrograde nature of eCB signaling that supports synapse-specific feedback can ensure the adaptive synaptic weakening only at active inputs. Concurrently, iSPNs associated with exploratory or alternative behaviors may become more active, providing the inhibitory bias required to suppress residual motor output and favour new action selection. Dopamine dips (reflecting a negative prediction error during omission)(72) may contribute by disinhibiting these iSPNs via reduced D2 receptor signaling. In overtrained mice, as well as in aged short-trained mice with rigid behavioural performance, impaired nanoscale remodeling of the mGluR_5_-eCB signaling complex during omission is equally important, because it prevents the appropriate silencing of iSPNs that are specifically tuned the outdated behavior. As a result, iSPNs associated with exploratory behaviors may continue to suppress alternative action plans, thereby maintaining the expression of the previously learned, but now suboptimal, response.

Central to this study is the focus on habit-related loss of behavioral flexibility, rather than behavioral flexibility *per se*, which we modeled by overtraining mice under a variable-interval schedule of reinforcement known to promote habit learning (67). Whether the molecular, cellular, and circuit plasticity mechanisms that are disrupted in association with the loss of behavioral flexibility in habitual mice overlap, at least in part, from those supporting flexibility in goal-directed behavior, remains an open question with potential implications for brain disorders or conditions characterized by an altered balance between cognitive/behavioral flexibility and rigidity. In this regard, aging increases reliance on antecedent stimulus–response (S–R) associations, leading to a greater tendency toward autonomous, habit-based behavioral control (8, 73–76). Our data indicate that short-trained, aging mice exhibited marked variability in their ability to update change in contingency, with some performing comparably to young, flexible mice and others resembling behaviorally inflexible, overtrained mice. The nanoscale upregulation of DAGLα and presynaptic CB_1_ receptors at cortico-iSPN synapses was observed only in aged mice performing in a flexible manner; these molecular changes were notably absent in their inflexible counterparts. These findings suggest that preserving retrograde eCB signaling plasticity in aging support behavioral adaptation, whereas its distruption contributes to inflexibility. In line with striatum-based mechanisms for age-related behavioral rigidity (77), rejuvenating striatal eCB functions (i.e., restoring mGlu_5_-DAGLα -CB_1_R coupling at cortico-DLS iSPN synapses) may help re-establishing adaptive control by suppressing the outdated action patterns.

## Materials and Methods

### Experimental procedures and experimental design

All procedures involving animals were carried out in accordance with the Italian Ministry of Health’s directives (D.lgs 26/2014) regulating animal research.

### Drugs

(RS)-3,5-DHPG, and SR 95531 hydrobromide (Gabazine), were purchased from Tocris Bioscience (Avonmouth, UK). Dilutions to final concentrations were made just before the start of each experiment in oxygenated aCSF.

### Animals

C57BL6J mice and drd2-EGFP (D_2_-EGFP) transgenic mice obtained from backcrossing C57BL6J and drd1-tdTomato/drd2-EGFP mice; at least 7–8 generations of backcrossing combined with systematic genotyping. BAC-transgenic mice (male, postnatal day – PND45-60) were housed in a controlled environment, on a 12 h light/dark cycle, with free access to water and/or food depending on each condition. Results may be specific to this sex and age range and may not be directly applicable to females or other developmental stages without further research.

### Behavioral experiments

#### Instrumental learning

Behavioral training and testing were performed in operant chambers (17.8 cm x 15.2 cm x 18.4 cm) housed within sound-attenuating chambers and equipped with two holes on either side of the food magazine (Med-associates, St Albans, VT). Mice were trained to nose-poke in one of two holes to obtain two reinforcers, either chocolate (F0 5301, Bilaney, UK) or sucrose pellets (F0 5684, Bilaney, UK). One reinforcer was delivered in the operant chamber contingent upon nose poking, into the magazine through a pellet dispenser. Magazine entries were recorded using an infrared beam. The reinforcer and nose poke hole were counterbalanced across groups. Before training, mice were food deprived, as to maintain 90% of their feeding body weight. Mice were fed daily after the training session. Initial nose-poke training consisted of two consecutive daily sessions of continuous reinforcement (CR), during which mice received a reinforcer for each nose poke. The session ended after 10 rewards. After the CR sessions, mice were trained on variable interval (VI) schedules (14, 16), in which active nose pokes were reinforced after variable time intervals that lasted on average 30 s (VI-30) or 60 s (VI-60), and ended after 20 reinforcers. After three daily VI-30 sessions, mice were either short-trained for four daily VI-60 sessions or overtrained for 18 VI-60 sessions (30)

#### Post-training omission procedure

The omission test started one day after instrumental training and lasted two days. On day 1, mice were exposed to a 30 min control session under the prevailing, positive action-outcome contingency (nose poking leads to reinforcer delivery). On day 2, mice were subjected to a 30 min session in which the previously learned A-O contingency was changed (negative contingency). That is, the pellet was delivered every 20 s without nose poke, but each nose poke would reset the counter and delay the food delivery. Thus, the new contingency does not require mice to learn a new behavior; instead, they must learn to withhold a behavior to maximize reward. The rates of active nose poke (ANP) under the two different A-O contingencies (negative A-O/positive A-O) were used to determine behavioral flexibility (29, 30, 66)

### Brain slice preparation

Brain slices containing the striatum and cortex were prepared as described (35, 78). Mice were anesthetized with isoflurane and decapitated, and their brains were transferred to ice-cold artificial cerebrospinal fluid (aCSF) containing 110 mM choline chloride, 2.5 mM KCl, 1.25 mM NaH2PO4, 7 mM MgCl2, 0.5 mM CaCl2, 25 mM NaHCO3, 25 mM D-glucose and 11.6 mM ascorbic acid, saturated with 95% O2 and 5% CO2 for dissection. Horizontal slices (270 µm thick) were prepared using a Vibratome 1000S slicer (Leica), then transferred to new aCSF containing 115 mM NaCl, 3.5 mM KCl, 1.2 mM NaH2PO4, 1.3 mM MgCl2, 2 mM CaCl2, 25 mM NaHCO3, and 25 mM D-glucose, and aerated with 95% O2 and 5% CO2. After incubating for 20 min at 32 °C, slices were kept at 22-24 °C. During subsequent experiments, slices were continuously superfused with aCSF at a rate of 2 ml/min at 28 °C.

### Electrophysiology

Whole-cell current-clamp recordings were performed in medium spiny projection neurons (SPNs) in the dorsolateral striatum in horizontal brain slices, which preserve intact cortico-striatal connections (79). Patch pipettes (4-6 MΩ) were filled with a solution containing 130 mM KMeSO4, 5 mM KCl, 5 mM NaCl, 10 mM HEPES, 2 mM MgCl2, 0.1 mM EGTA, 0.05 mM CaCl2, 2 mM Na2ATP and 0.4 mM Na3GTP (pH 7.2-7.3, 280-290 mOsm/kg). After the recording, neurobiotin (0.5 mg/ml, Vector Laboratories,Peterborough, United Kingdom) was added to the pipette solution to identify SPNs of the indirect (iSPN) and direct (dSPN) pathways, by immunolabeling for the A2A receptor (marker of iSPNs) and substance P (marker of dSPNs) as described before (35, 78). During recordings, SPNs were clamped at a holding membrane potential of -60 mV. Excitatory postsynaptic potentials (EPSPs) were evoked in presence of the GABAA receptor antagonist gabazine (10 µM bath applied) by electrical stimulation of the somatosensory cortex layer 5, using a concentric bipolar electrode (40-100 μA; 40-80 μs; FHC, Bowdoin, Maine), and acquired every 10 s. After 10 minutes recording the inhibition of EPSP amplitude was induced by bath application of the mGlu_1/5_ agonist (RS)-3,5-DHPG (100 μM). 15 minutes after DHPG application, evoked EPSCs were recorded in presence of gabazine for the remaining time. Data were acquired using a Multiclamp 700B amplifier controlled by pClamp 10 software (Molecular Device), filtered at 10 kHz and sampled at 20 kHz (Digidata 1322, Molecular Device). All data are reported without corrections for liquid junction potentials. Data where the input resistance (Rinp) changed >20% have been excluded. The occurrence and magnitude of synaptic plasticity was evaluated by comparing the normalized EPSP amplitudes from the last 5 minutes of baseline recordings with the corresponding values at 30–35 minutes after DHPG application. LTD plots were generated by averaging the peak amplitude of individual EPSPs in 2-min bins. Data were analyzed by one-way repeated-measures ANOVA (RM1W) for comparisons within a group (GraphPad Instat 3 software). Parametric statistics were utilized for comparisons of two groups (independent two-sample t test), unless data were not normally distributed (Mann-Whitney-U-Test) (GraphPad Instat 3 software).

### Correlated confocal and STORM microscopy

#### Sample preparation

D_2_-EGFP transgenic mice were anesthetized under isofluorane, perfused transcardially with 0.9% (m/v) saline for 2 min, followed by 4% PFA in 0.1 M phosphate buffer (PB, pH 7.4) for 20 min at a rate of 5 mL/min. Brains were then postfixed 2 hours at room temperature in PFA 4% PB 0.1M; washed in PB 0.1M; and stored at +4°C in 0.1M PB supplemented with sodium azide 0.05%. The 20-μm-thick coronal sections were cut using a Leica VT-1000S vibratome.

#### Immunolabeling

After slicing, the striatal sections were immunolabeled in a free-floating manner. Following washing in 0.1M PB, and 0.05 M Tris-buffered saline (TBS), the sections were blocked with 5% (m/v) NDS (normal donkey serum, Sigma) in TBS supplemented with 0.3% (v/v) Triton X-100 for 45 minutes. The sections were then incubated with affinity-purified primary antibodies in TBS overnight at room temperature (20–25 °C) on an orbital shaker. Primary antibodies used in this study included rabbit antibody against DAGLα (used in 1:1000, a kind gift from Ken Mackie)(80), guinea pig antibody against DAGLα (1:1000, gift from M. Watanabe. Epitope: C-terminal 42 Aa)(81), rabbit antibody against CB_1_ receptor (1:1000, a kind gift from Ken Mackie); rabbit antibody against mGlu_5_ (1:1000, Sigma-Aldrich AB5675); goat antibody against VGLUT1 (1:1000, a kind gift from Masahiko Watanabe)(82); goat antibody against EGFP (1:1000, EGFP goat: Abcam, catalog number: ab5450). The specificity of the antibodies were verified in the original publications by using knockout mice (80–83). Sections were then washed in TBS and incubated with an Alexa Fluor 488-conjugated anti-goat (1:400, Jackson, 705-545-147) secondary antibody, along with the appropriate STORM-compatible secondary antibodies (anti-rabbit CF568, Biotium, 20098-1, 1:1000; anti-rabbit Alexa Fluor 647, Jackson, 711-605-152, 1:400; anti-guinea pig Alexa Fluor 647, Jackson, 706-605-148, 1:400). Free-floating sections were then washed in TBS two times 20 minutes, and in PB two times 10 minutes. These steps were followed by 4% PFA post-fixation for 10 minutes, followed by three washes of 10 minutes in PB. Sections were mounted and dried on acetone-cleaned #1.5 borosilicate coverslips, and stored dry at 4 °C until imaging.

#### Imaging

Samples were covered with imaging medium freshly prepared as described previously (47) containing 5% (m/v) glucose, 0.1 M 2-Mercaptoethylamine (MEA, also known as cysteamine, Sigma-Aldrich, cat. no. 30070-10G), 1 mg ml^−1^ glucose oxidase and catalase (2.5 μl ml^−1^ of aqueous solution from Sigma, approximately 1,500 U ml^−1^ final concentration) in Dulbecco’s PBS (Sigma) right before the start of the imaging process. Abbelight Smart Kit Buffer was used for the dual STORM imaging of mGlu_5_ and DAGLα. Coverslips were sealed with nail polish, and transferred into the microscope setup after a 10 min drying period. STORM imaging was performed for up to 1h after covering the specimens.

The STORM images and the correlated high-power confocal stacks were acquired via a CFI Apo TIRF 100× objective (1.49 NA) on a Nikon Ti-E inverted microscope equipped with a Nikon N-STORM system, a Nikon C2 confocal scan head and an Andor iXon Ultra 897 EMCCD. The setup was controlled by Nikon NIS-Elements AR software with N-STORM module. For STORM imaging, a 300-mW laser was used (VFL-P-300-647, MPB Communications), fiber-coupled to the laser board of the microscope setup. To obtain images, the field of view was selected by using the live EMCCD image under 488-nm illumination. A confocal z-stack (512 × 512 × 15 voxels, 80 × 80 × 150 nm resolution in x,y,z dimensions, respectively) was then acquired. The field of view of the confocal scan area and the EMCCD were set to be identical. For increased consistency, the specimen was illuminated with maximal 647 imaging laser power (>2 kW/cm^2^ in the field of view) until the image was no longer saturated—i.e., most of the fluorophores were kept in the ‘off’ state (1–2 s) before starting the STORM imaging. The STORM image was acquired using a STORM filter cube (Nikon) and the EMCCD STORM image was acquired using a far-red STORM filter cube (Nikon). Altogether 10,000 frames with 30 ms exposure times were recorded. A 10 % 405 nm activation laser power aided the continuous reactivation of the flurophores. Dual-color STORM imaging was performed in a sequential manner, starting as above and followed by another 5,000 frames using 561 nm laser excitation and a HQ Red filter cube. To minimize out-of-focus background, oblique illumination was applied using the TIRF illuminator of the microscope. Localization points were collected at a similar tissue depth (∼5 μm) for all images to equalize antibody penetration probabilities and the effect of light scattering.

#### Image processing

Confocal images were deconvolved to reduce the effect of light scattering on overestimating profile sizes as confocal stacks provide structural information for the super-resolution image by using the classic maximum likelihood estimation (CMLE) algorithm in the Huygens Professional software (SVI). The refractive index of the imaging medium is considered to be equal to water (1.338). Stacks were converted to OME (.ome.tif) image format after deconvolution.

Peak detection on STORM images was run through the NIS-Elements N-STORM module, which is based on the 3D-DAOSTORM algorithm (84, 85). 3D calibration data was obtained by imaging 100 nm large fluorescent beads and the measurements were loaded to extract 3D coordinates. The peak intensity threshold was set at 1,000 grey levels to avoid noise detection. Chromatic aberrations were corrected by using individual *z*-calibration curves for each wavelength, and by warp calibration. After peak detection analysis, each data file was exported in text format as a molecule list. NIS-Elements STORM module also performed image-based correction for sample drift in *x*, *y* and *z*.

#### Correlated confocal and STORM image analysis

Deconvolved confocal images and STORM localization points were analyzed by the VividSTORM software (47). The filtering of DAGLα, mGlu_5_ or CB_1_ STORM localization points (LPs) based on EGFP and VGLUT1-immunolabeling was carried out using VividSTORM. First, the correlated confocal and STORM images were aligned in VividSTORM. Next, the region-of-interest (ROI) containing an EGFP-expressing spine head adjacent to a VGLUT1-positive cortico-striatal axon terminal was selected. The target profile was outlined using VividSTORM active contour selection of the appropriate channel (EGFP for mGlu_5_ and DAGLα, VGLUT1 for CB_1_). In each ROI, the number of localization points per channels and their coordinates were exported. Clustering analysis was carried out using DBSCAN analysis integrated into VividSTORM. A cluster was defined by at least 5 LPs within 60 nm distances of each other. For DAGLα - mGlu_5_ LP nearest neighbor distance measurements, the minimum Euclidean distance between DAGLα LPs and the mGlu_5_ LP was measured.

### Statistical analyses

Appropriate parametric statistics were used to test hypotheses, unless data did not meet the assumptions of the intended parametric test (normality test). In this case, appropriate non-parametric tests were used. Power analysis specifications to estimate sample size were: power = 0.8, alpha = 0.05, two-tailed, and an effect size that is 50% greater than previously observed standard deviations. Data were analyzed by two-way repeated measure ANOVA or one-way repeated measure ANOVA for comparisons within a group, and one-way ANOVA (1WA) for between-group comparisons (GraphPad Prism 9 software). A mixed-effects two-way ANOVA was used to analyze experiments with between-subjects (short-and overtraining) and within-subjects variables (post-training positive A-O versus negative A-O or valued versus devalued conditions). Corrected post-hoc tests (Tukey or Sidak as indicated) were performed only when the ANOVA yielded a significant main or interaction effect. Two groups were tested for statistical significance using the independent samples t-test, the paired samples t-test, or equivalent non-parametric tests (GraphPad Prism 9 software). Behavioral analysis using the K-Nearest Neighbors (KNN) algorithm was performed with a custom Python script (86), which can be substituted with Python’s scikit-learn KNeighborsClassifier (87) setting k to 8. STORM NLP and clusters were analyzed using Mann-Whitney tests.

Statistical details of the experiments are shown in the Results, Figure legends, and are summarized in **Table 1**.

## Author contributions (CrediT – Contributor Roles Taxonomy)

**Conceptualization**: V.PB., I.K., R.T.

**Methodology**: V.PB., M.Z., A.C., I.K., R.T.

**Investigation**: V.PB., A.C.

**Formal Analysis:** V.PB., M.Z., A.C., V.M., B.D.

**Software:** V.M., B.D.

**Writing – Original Draft:** V.PB., M.Z., I.K., R.T.

**Writing – Review & Editing:** I.K., R.T.

**Supervision:** I.K., R.T

**Project Administration:** I.K., R.T.

**Funding Acquisition:** I.K., R.T.

## Supporting information

Statistical Table

## Acknowledgments

The authors are grateful to Alice Gino, E. Tischler, B. Pintér, J.K. Leffel for their technical support. The authors also appreciate the support of the Nikon Microscopy Center at the Institute of Experimental Medicine, Nikon Europe B.V., Nikon Austria GmbH, and Auro-Science Consulting. IK holds the Naus Family Chair in Addiction Sciences in the Department of Psychological and Brain Sciences at Indiana University Bloomington. This work was partially funded by the Fondazione Istituto Italiano di Tecnologia (RT), by the project “National Center for Gene Therapy and Drug based on RNA Technology” (CN00000041) and NextGenerationEU PNRR MUR – M4C2 – Action 1.4-Call “Potenziamento strutture di ricerca e di campioni nazionali di R&S” (CUP J33C22001130001) (RT), and has been carried out within the IIT-RNA Flagship Programme, by the NKFIH EXCELLENCE program 151377 (IK) and the National Institutes of Health grant P30DA056410 (IK).

---

**Figure S1, related to Figure 1.**

(**A**) Schematic depicts the behavioral paradigm, CR: continuous reinforcement, VI: variable interval 30 s (VI-30) or 60 s (VI-60).

(**B**) Averaged time courses of magazine entry (ME) rates (left) and inactive nose-poke (INP) rates (right) during training in C57Bl6J mice (Sh, n=23; Ov, n=22; Sh ΔA-O, n=17, Ov ΔA-O, n=18; ME/min, session: F_8,592_ = 40.31, ****p < 0.0001, group: F_3,74_ = 0.7964, p = 0.5, interaction: F_24,592_ = 1.289, p = 0.16; IN/min, session: F_8,592_ = 5.186, ****p < 0.0001, group: F_3,74_ = 0.6303, p = 0.5978, interaction: F_24,592_ = 1.955, **p = 0.0045).

(**C**) INP rates during the post-training omission procedure in short-and overtrained C57Bl6J mice (Sh n = 17, Ov n = 16; A-O contingency: F_1,31_ = 0.1839, p = 0.67, group: F_1,31_ = 0.09785 p = 0.7565, A-O contingency X group interaction: F_1,31_ = 1.422, p = 0.2421; Sh: positive A-O: 0.41 ± 0.16, negative A-O, 0.29 ± 0.05, Sidak: p = 0.4932; Ov: positive A-O: 0.37 ± 0.08, negative A-O: 0.44 ± 0.09, Sidak: p = 0.7925). All data are expressed as mean ± SEM.

(**D**) mGlu-LTD at cortico-dSPN synapses. In dSPNs, DPHG application induced an LTD of EPSPs of similar magnitude across all training regimes (Sh: n = 9 cells, N = 9 mice, 80.3 ± 5.8%; ShΔAO: n = 9 cells, N = 8 mice; 78.6 ± 2.8%; Ov: n = 9 cells, N = 7 mice, 87.2 ± 6.9; OvΔAO: n = 12 cells, N = 12 mice, 84.7 ± 3.; p = 0.6530).

(**E**) ANP rates during instrumental conditioning in short-and overtrained mice (Sh, n = 14; Ov, n = 14), and in short-and overtrained mice subjected to changes in contingency (Sh ΔA-O, n = 16, Ov ΔA-O, n = 11) in the D2-EGFP transgenic mouse line.

(**F**) (Left) Post-training omission procedure in D_2_-EGFP mice. Comparison of ANP rates between positive and negative A-O contingency in both short- and overtrained mice (Sh n = 16, positive A-O: 18.71 ± 1.91, negative A-O: 11.75 ± 1.69, Sidak ****p < 0.0001; Ov n = 11, positive A-O: 19.66 ± 2.57, negative A-O 20.08 ± 2.16; Sidak p = 0.9483). (Right) Time course of ANP ratios (ANP rates under negative A-O/ANP rates under positive A-O) in short- and overtrained D_2_-EGFP mice.

(**G**) Number of obtained reinforcers (pellets; g) in the positive- and negative A-O contingency sessions (Sh n = 16, positive A-O: 0.38 ± 0.01, Ov n = 11, positive A-O: 0.39 ± 0.01, p = 0.9736; Sh negative A-O: 0.29 ± 0.05; Ov negative A-O 0.11 ± 0.04; Sidak ****p < 0.0001).

(**H**) Averaged time courses of magazine entry (ME) rates (left) and inactive nose-poke (INP) rates (right) during training in D_2_–EGFP mice (Sh, n=12; Ov, n=14; Sh ΔA-O, n=16, Ov ΔA-O, n=11; ME/min, session: F_8, 392_ = 6.569, ****p < 0.0001, group: F_3, 49_ = 0.7830, p = 0.5092, interaction: F_24, 392_ = 1.121, p = 0.3167; IN/min, session: F_8, 392_ = 2.308, *p = 0.0199, group: F_3, 49_ = 0.5313, p = 0.663, interaction: F_24, 392_ = 1.680, *p = 0.0246).

(**I**) INP rates during the post-training omission procedure in short- and overtrained D_2_–EGFP mice (Sh n = 16, Ov n = 11; A-O contingency: F_1,25_ = 2.954, p = 0.098, group: F_1,25_ = 1.472 p = 0.2364, A-O contingency X group interaction: F_1,25_ =4.424, *p = 0.457; Sh: positive A-O: 0.18 ± 0.04, negative A-O, 0.17 ± 0.03,: p = 0.9451; Ov: positive A-O: 0.22 ± 0.08, negative A-O: 0.34 ± 0.11, Sidak: *p = 0.0398). All data are expressed as mean ± SEM.

**Figure S2, related to Figure 4.**

(**A**) Schematic of the behavioral paradigms in the three cohorts (Sh; Sh_Aged; Sh_Aged ΔA-O) , CR: continuous reinforcement, VI: variable interval 30 s (VI-30) or 60 s (VI-60).

(**B**) Averaged time courses of magazine entry (ME) rates (left) and inactive nose-poke (INP) rates (right) during training (Sh, n = 10; Sh_Aged, n = 10; Sh_Aged ΔA-O, n = 17; ME/min, session: F_8, 272_ = 7.967, ****p < 0.0001, group: F_2, 34_ = 0.6414, p = 0.5328, interaction: F_16, 272_= 0.4597, p = 0.9638; IN/min, session: F_8, 272_ = 4.535, ****p < 0.0001, group: F_2, 34_ = 0.4036, p = 0.6711, interaction: F_16, 272_ = 1.377, p = 0.1525).

(**C**) (Left) Post-training omission procedure in aged short-trained mice (Sh_Aged ΔAO Flexible n = 10, Sh_Aged ΔAO Inflexible n = 7). INP rates (A-O contingency: F_1, 15_ = 4.837, *p = 0.0440, group: F_1, 15_ = 3.145, p = 0.0964, A-O contingency X group interaction: F_1, 15_ = 5.118, *p = 0.0390; Sh_Aged ΔAO Flexible: positive A-O: 0.14 ± 0.03, negative A-O, 0.11 ± 0.02, Sidak: p = 0.9985; Sh_Aged ΔAO Inflexible: positive A-O: 0.14 ± 0.05, negative A-O: 0.58 ± 0.25, Sidak: *p = 0.0215. All data are expressed as mean ± SEM.

**Table 1:** Statistical table

## Notes

### Competing Interest Statement

The authors have declared no competing interest.

