## Supplementary figures and images for "Nanoscale synaptic remodeling at corticostriatal circuits predicts flexible action control"

### Supplemental Figure 1

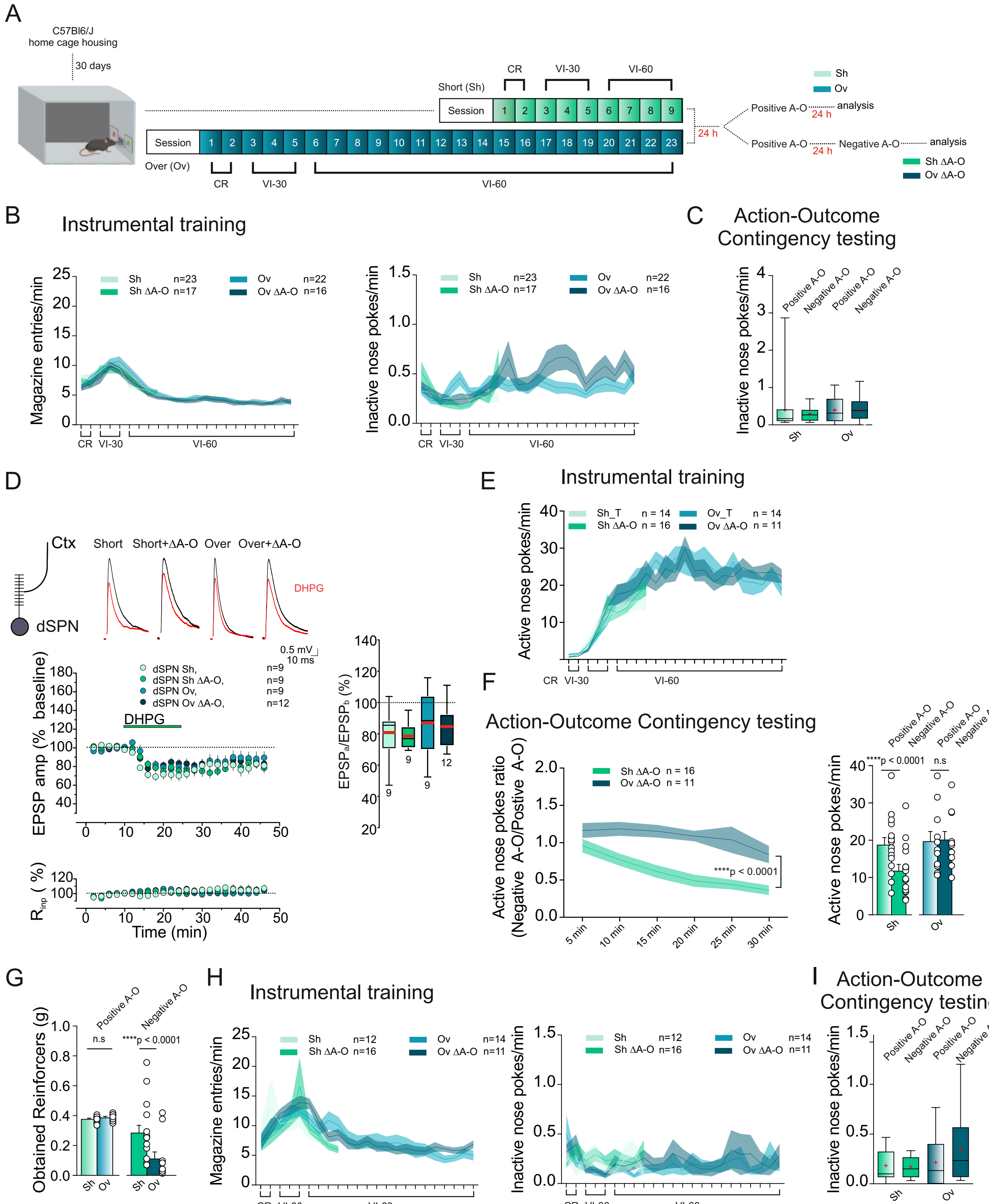

### Supplemental Figure 2

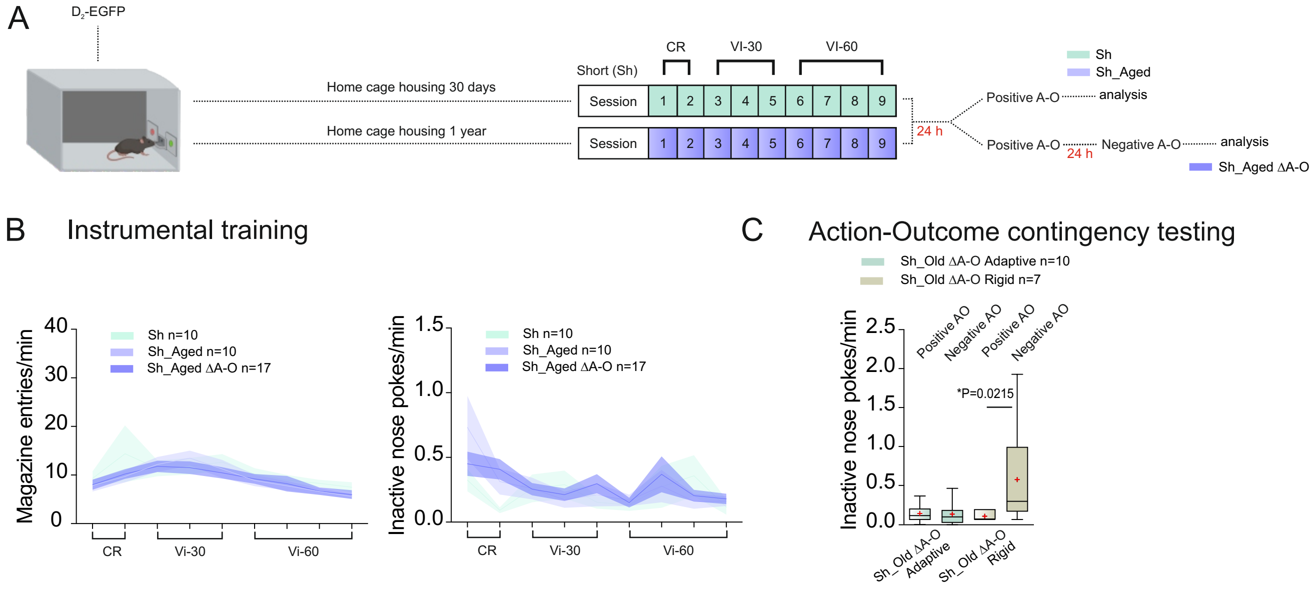
