## Supplementary material for "Nanoscale synaptic remodeling at corticostriatal circuits predicts flexible action control": Statistical Table

### Table 1

| Figure | Variable | Condition | Mean ± SEM. n = | Source of Variation | F (DFn. DFd) | p value | Post-Hoc (Šídák) | P value summary | Test |
|---|---|---|---|---|---|---|---|---|---|
| **Fig. 1B** | ANPmin | | | | | | | | Two-way RM ANOVA |
| | Sh | | n = 23 | Session x Group | F (24. 592) = 1.044 | P=0.4058 | | | |
| | ShΔAO | | n = 17 | Session | F (8. 592) = 167.2 | ****P<0.0001 | | | |
| | Ov | | n = 22 | Group | F (3. 74) = 1.003 | P=0.3963 | | | |
| | OvΔAO | | n = 16 | | | | | | |
| **Fig. 1C** | ANPmin | | | | | | | | Two-way RM ANOVA |
| | ShΔAO | 5 min | 0.93 ± 0.06. n = 17 | Time x Group | F (5. 155) = 1.147 | P=0.3382 | | | |
| | | 10 min | 0.88 ± 0.08. n = 17 | Time | F (5. 155) = 10.35 | ****P<0.0001 | | | |
| | | 15 min | 0.72 ± 0.09. n = 17 | Group | F (1. 31) = 5.067 | *P=0.0316 | | | |
| | | 20 min | 0.59 ± 0.08. n = 17 | | | | | | |
| | | 25 min | 0.50 ± 0.06. n = 17 | | | | | | |
| | | 30 min | 0.39 ± 0.06. n = 17 | | | | | | |
| | OvΔAO | 5 min | 1.12 ± 0.15. n = 16 | | | | | | |
| | | 10 min | 1.01 ± 0.11. n = 16 | | | | | | |
| | | 15 min | 0.91 ± 0.11. n = 16 | | | | | | |
| | | 20 min | 0.94 ± 0.13. n = 16 | | | | | | |
| | | 25 min | 0.74 ± 0.12. n = 16 | | | | | | |
| | | 30 min | 0.81 ± 0.16. n = 16 | | | | | | |
| | ANPmin | | | | | | | | |
| | ShΔAO | Positive AO | 13.40 ± 1.29. n = 17 | Group x A-O Contingency | F (1. 31) = 1.512 | P=0.2280 | Sh_Positibve AO vs. Sh_Negative AO ***P=0.0009 | | MD Anova |
| | | Negative AO | 9.05 ± 1.12. n = 17 | Group | F (1. 31) = 2.436 | P=0.1287 | | | |
| | | | | A-O Contingency | F (1. 31) = 17.74 | ***P=0.0002 | Ov_ Positive AO vs. Ov_ Negative AO P=0.0902 | | |
| | OvΔAO | Positive AO | 14.76 ± 1.21. n = 16 | | | | | | |
| | | Negative AO | 12.37 ± 1.17. n = 16 | | | | | | |
| | Obtained reinforcers | | | | | | | | |
| | ShΔAO | Positive AO | 0.38 ± 0.04. n = 17 | A-O Contingency x Group | F (1. 35) = 10.53 | **P=0.0026 | Sh_ Positive AO vs. Ov_ Positive AO P=0.9850 | | MD Anova |
| | OvΔAO | Positive AO | 0.39 ± 0.02. n = 16 | A-O Contingency | F (1. 35) = 21.82 | ****P<0.0001 | | | |
| | | | | Group | F (1. 35) = 14.76 | ***P=0.0005 | Sh_ Negative AO vs. Ov_ Negative AO ****P<0.0001 | | |
| | ShΔAO | Negative AO | 0.34 ± 0.21. n = 17 | | | | | | |
| | OvΔAO | Negative AO | 0.17 ± 0.10. n = 16 | | | | | | |
| **Fig. 1D** | Sh | | 77,76 ± 6,090 n = 5 | Group | F (3, 23) = 5,207 | **P = 0,0068 | Sh vs Sh_Negative AO | *P<0.05 | One-way ANOVA |
| | ShΔAO | | 50,79 ± 8,722 n = 5 | | | | Ov vs Sh_Negative AO | *P<0.05 | |
| | Ov | | 75,11 ± 5,465 n = 8 | | | | Sh_Negative AO vs Ov_Negative AO | **P<0.01 | |
| | OvΔAO | | 84,44 ± 4,700 n = 9 | | | | Dunnett's multiple comparisons test | | |
| **Fig. 1E** | mGLU5 LP / µm2 | | | | | | Dunn's multiple comparisons test | | Kruskal-Wallis test |
| | | Sh | 1.00 ± 0.09 n = 89 | | | ****p<0.0001 | Sh vs. ShΔAO ****P=<0.0001 | | |
| | | ShΔAO | 2.27 ± 0.13 n = 109 | | | | Sh vs. Ov P=0.9567 | | |
| | | Ov | 1.25 ± 0.12 n = 80 | | | | Sh vs. OvΔAO P=>0.9999 | | |
| | | OvΔAO | 1.15 ± 0.10 n = 68 | | | | ShΔAO vs. Ov ****P=<0.0001 | | |
| | | | | | | | ShΔAO vs. OvΔAO ****P=<0.0001 | | |
| | | | | | | | Ov vs. OvΔAO P=>0.9999 | | |
| | mGLU5 Clusters / µm2 | | | | | | Dunn's multiple comparisons test | | Kruskal-Wallis test |
| | | Sh | 1.00 ± 0.20 n = 89 | | | p=0,1837 | Sh vs. ShΔAO P=0.4767 | | |
| | | ShΔAO | 1.24 ± 0.18 n = 109 | | | | Sh vs. Ov P=>0.9999 | | |
| | | Ov | 1.14 ± 0.25 n = 80 | | | | Sh vs. OvΔAO P=>0.9999 | | |
| | | OvΔAO | 1.16 ± 0.23 n = 68 | | | | ShΔAO vs. Ov P=0.3161 | | |
| | | | | | | | ShΔAO vs. OvΔAO P=>0.9999 | | |
| | | | | | | | Ov vs. OvΔAO P=>0.9999 | | |
| | mGLU5 LP / Clusters | | | | | | Dunn's multiple comparisons test | | Kruskal-Wallis test |
| | | Sh | 1.00 ± 0.11 n = 27 | | | p=0,5528 | P=>0.9999 | | |
| | | ShΔAO | 1.55 ± 0.25. n = 51 | | | | Sh vs. Ov P=>0.9999 | | |
| | | Ov | 0.85 ± 0.07. n = 21 | | | | Sh vs. OvΔAO P=>0.9999 | | |
| | | OvΔAO | 1.31 ± 0.39. n = 20 | | | | ShΔAO vs. Ov P=>0.9999 | | |
| | | | | | | | ShΔAO vs. OvΔAO P=>0.9999 | | |
| | | | | | | | Ov vs. OvΔAO P=>0.9999 | | |
| **Fig. 2B** | DAGLα LP / µm2 | | | | | | Dunn's multiple comparisons test | | Kruskal-Wallis test |
| | | Sh | 1 ± 0.11 n = 116 | | | ****p<0.0001 | Sh vs. Sh ΔAO ****P=<0.0001 | | |
| | | ShΔAO | 3.03 ± 0.28 n = 143 | | | | Sh vs. Ov ***P=0.0009 | | |
| | | Ov | 0.54 ± 0.07 n = 149 | | | | Sh vs. Ov ΔAO P=0.9014 | | |
| | | OvΔAO | 0.78 ± 0.11 n = 109 | | | | Sh ΔAO vs. Ov ****P=<0.0001 | | |
| | | | | | | | Sh ΔAO vs. Ov ΔAO ****P=<0.0001 | | |
| | | | | | | | Ov vs. Ov ΔAO P=0.1698 | | |
| | DAGLα Clusters / µm2 | | | | | | Dunn's multiple comparisons test | | Kruskal-Wallis test |
| | | Sh | 1 ± 0.18 n = 116 | | | ****p<0.0001 | Sh vs. Sh ΔAO ****P=<0.0001 | | |
| | | ShΔAO | 2.39 ± 0.27 n = 143 | | | | Sh vs. Ov **P=0.0034 | | |
| | | Ov | 0.34 ± 0.1 n = 149 | | | | Sh vs. Ov ΔAO *P=0.0354 | | |
| | | OvΔAO | 0.4 ± 0.12 n = 109 | | | | Sh ΔAO vs. Ov ****P=<0.0001 | | |
| | | | | | | | Sh ΔAO vs. Ov ΔAO ****P=<0.0001 | | |

| | | | | | | | | |
|---|---|---|---|---|---|---|---|---|
| | | | | | | | Ov vs. Ov ΔAO P=>0.9999 | |
| | DAGLα LP / Clusters | | | | | | Dunn's multiple comparisons test | Kruskal-Wallis test |
| | | Sh | 1 ± 0.06 n = 36 | | | ***p=0,0003 | Sh vs. Sh ΔAO **P=0.0053 | |
| | | ShΔAO | 1.76 ± 0.15 n = 90 | | | | Sh vs. Ov P=>0.9999 | |
| | | Ov | 1.01 ± 0.12 n = 16 | | | | Sh vs. Ov ΔAO P=>0.9999 | |
| | | OvΔAO | 1.06 ± 0.17 n = 15 | | | | Sh ΔAO vs. Ov *P=0.0491 | |
| | | | | | | | Sh ΔAO vs. Ov ΔAO *P=0.0426 | |
| | | | | | | | Ov vs. Ov ΔAO P=>0.9999 | |
| **Fig. 2C** | Minimum Distance (nm) DGL LP - mGLUR5 LP | | | | | | Dunn's multiple comparisons test | Kruskal-Wallis test |
| | | Sh | 135.80 ± 8.80 n = 88 | | | ****p<0.0001 | Sh vs. ShΔAO ***P=0.0002 | |
| | | ShΔAO | 91.43 ± 6.12 n = 115 | | | | Sh vs. Ov P=0.6677 | |
| | | Ov | 156.90 ± 10.24 n = 73 | | | | Sh vs. OvΔAO P=0.9609 | |
| | | OvΔAO | 147.90 ± 8.78 n = 67 | | | | ShΔAO vs. Ov ****P=<0.0001 | |
| | | | | | | | ShΔAO vs. OvΔAO ****P=<0.0001 | |
| | | | | | | | Ov vs. OvΔAO P=>0.9999 | |
| **Fig. 3B** | Cb1 LP / μm2 | | | | | | Dunn's multiple comparisons test | Kruskal-Wallis test |
| | | Sh | 1 ± 0.21 n = 71 | | | ****p<0.0001 | Sh vs. Sh ΔAO ****P=<0.0001 | |
| | | ShΔAO | 3.79 ± 0.49 n = 97 | | | | Sh vs. Ov P=>0.9999 | |
| | | Ov | 1.02 ± 0.11 n = 100 | | | | Sh vs. Ov ΔAO P=>0.9999 | |
| | | OvΔAO | 1.28 ± 0.23 n = 72 | | | | Sh ΔAO vs. Ov ****P=<0.0001 | |
| | | | | | | | Sh ΔAO vs. Ov ΔAO ****P=<0.0001 | |
| | | | | | | | Ov vs. Ov ΔAO P=>0.9999 | |
| | Cb1 Clusters / μm2 | | | | | | Dunn's multiple comparisons test | Kruskal-Wallis test |
| | | Sh | 1 ± 0.3 n = 71 | | | ****p<0.0001 | Sh vs. Sh ΔAO ****P=<0.0001 | |
| | | ShΔAO | 4.87 ± 0.74 n = 97 | | | | Sh vs. Ov P=>0.9999 | |
| | | Ov | 1.16 ± 0.27 n = 100 | | | | Sh vs. Ov ΔAO P=>0.9999 | |
| | | OvΔAO | 1.61 ± 0.42 n = 72 | | | | Sh ΔAO vs. Ov ****P=<0.0001 | |
| | | | | | | | Sh ΔAO vs. Ov ΔAO ****P=<0.0001 | |
| | | | | | | | Ov vs. Ov ΔAO P=>0.9999 | |
| | Cb1 LP / Clusters | | | | | | Dunn's multiple comparisons test | Kruskal-Wallis test |
| | | Sh | 1 ± 0.14 n = 15 | | | p=0,1019 | Sh vs. Sh ΔAO P=>0.9999 | |
| | | ShΔAO | 1.16 ± 0.11 n = 56 | | | | Sh vs. Ov P=0.3798 | |
| | | Ov | 0.73 ± 0.07 n = 19 | | | | Sh vs. Ov ΔAO P=>0.9999 | |
| | | OvΔAO | 1.06 ± 0.23 n = 17 | | | | Sh ΔAO vs. Ov P=0.128 | |
| | | | | | | | Sh ΔAO vs. Ov ΔAO P=>0.9999 | |
| | | | | | | | Ov vs. Ov ΔAO P=>0.9999 | |
| **Fig. 4B** | | | | | | | | |
| Adaptive Index / MGlu5 LP/μm2 | | | | | ***P = 0.0004 | | | |
| | | | | | r2 = 0,8967 | | | |
| Adaptive Index / mGlu5 Clusters/μm2 | | | | | P = 0,3754 | | | |
| | | | | | r2 = 0,1325 | | | |
| Adaptive Index / mGlu5 LP / Cluster | | | | | P = 0,4456 | | | |
| | | | | | r2 = 0,09991 | | | |
| **Fig. 4C** | | | | | | | | |
| Adaptive Index / DGL LP/μm2 | | | | | **P = 0,0015 | | | |
| | | | | | r2 = 0,8875 | | | |
| Adaptive Index / DGL Clusters/μm2 | | | | | **P = 0.0040 | | | |
| | | | | | r2 = 0.8348 | | | |
| Adaptive Index / DGL LP / Cluster | | | | | P = 0,3908 | | | |
| | | | | | r2 = 0,1499 | | | |
| **Fig. 4D** | | | | | | | | |
| Adaptive Index / Cb1 LP/μm2 | | | | | **P = 0,0026 | | | |
| | | | | | r2 = 0,8613 | | | |
| Adaptive Index / Cb1 Clusters/μm2 | | | | | *P = 0,0244 | | | |
| | | | | | r2 = 0,6698 | | | |
| Adaptive Index / Cb1 LP / Cluster | | | | | P = 0,6048 | | | |
| | | | | | r2 = 0,05740 | | | |
| **Fig. 5B** | | | | | | | | Two-way RM ANOVA |
| | ANPmin | | | | | | | |
| | Sh | n = 10 | | Session x Group | F (16, 272) = 0.7876 | P=0.6994 | | |
| | Sh_Aged | n = 10 | | Session | F (8, 272) = 75.42 | ****P<0.0001 | | |
| | Sh_Aged ΔAO | n = 17 | | Group | F (2, 34) = 0.07062 | P=0.9320 | | |

| | | | | | | | | | |
|---|---|---|---|---|---|---|---|---|---|
| **Fig. 5D** | ANPmin | | | | | | | | Two-way RM ANOVA |
| | Sh_Aged ΔAO Flexil | 5 min | 1.07 ± 0.10. n = 10 | Time x Group | F (5. 75) = 8.983 | ****P<0.0001 | | | |
| | | 10 min | 0.88 ± 0.16. n = 10 | Time | F (5. 75) = 17.41 | ****P<0.0001 | | | |
| | | 15 min | 0.47 ± 0.11. n = 10 | Group | F (1. 15) = 37.69 | ****P<0.0001 | | | |
| | | 20 min | 0.19 ± 0.05. n = 10 | | | | | | |
| | | 25 min | 0.09 ± 0.03. n = 10 | | | | | | |
| | | 30 min | 0.07 ± 0.04. n = 10 | | | | | | |
| | Sh_Aged ΔAO Inflex | 5 min | 1.13 ± 0.12. n = 7 | | | | | | |
| | | 10 min | 0.82 ± 0.03. n = 7 | | | | | | |
| | | 15 min | 0.91 ± 0.11. n = 7 | | | | | | |
| | | 20 min | 0.69 ± 0.06. n = 7 | | | | | | |
| | | 25 min | 0.89 ± 0.06. n = 7 | | | | | | |
| | | 30 min | 0.91 ± 0.11. n = 7 | | | | | | |
| | ANPmin | | | | | | | | MD Anova |
| | Sh_Flexible ΔAO | Positive AO | 12.05 ± 2.13. n = 10 | Group x Condition | F (1. 15) = 3.437 | P=0.0835 | | | |
| | | Negative AO | 5.89 ± 1.13. n = 10 | Group | F (1. 15) = 18.23 | ***P=0.0007 | Control - Reversal | | |
| | | | | Condition | F (1. 15) = 22.97 | ***P=0.0002 | Short GD | ***P=0.0002 | |
| | Sh_Inflexible ΔAO | Positive AO | 22.20 ± 2.46. n = 7 | | | | Short Hab | P=0.1437 | |
| | | Negative AO | 19.48 ± 2.60. n = 7 | | | | | | |
| **Fig. 5E** | DAGLα LP / μm2 | | | | | | Dunn's multiple comparisons test | | Kruskal-Wallis test |
| | | Sh | 1 ± 0.18 n = 126 | | | ****p<0.0001 | Sh vs. Sh Old P=>0.9999 | | |
| | | Sh Aged | 0.85 ± 0.13 n = 162 | | | | Sh vs. Sh Old ΔAO Flexible ***P=0.0002 | | |
| | | Sh Aged ΔAO Flexibl | 2.22 ± 0.44 n = 113 | | | | Sh vs. Sh Old ΔAO Inflexible *P=0.0246 | | |
| | | Sh Aged ΔAO Inflexib | 0.56 ± 0.11 n = 131 | | | | Sh Old vs. Sh Old ΔAO Flexible ****P=<0.0001 | | |
| | | | | | | | Sh Old vs. Sh Old ΔAO Inflexible *P=0.0307 | | |
| | | | | | | | Sh Old ΔAO Flexible vs. Sh Old ΔAO Inflexible ****P=<0.0001 | | |
| | DAGLα Clusters / μm2 | | | | | | Dunn's multiple comparisons test | | Kruskal-Wallis test |
| | | Sh | 1 ± 0.31 n = 126 | | | ****p<0.0001 | Sh vs. Sh Old P=>0.9999 | | |
| | | Sh Aged | 0.66 ± 0.18 n = 162 | | | | Sh vs. Sh Old ΔAO Flexible ****P=<0.0001 | | |
| | | Sh Aged ΔAO Flexibl | 1.89 ± 0.28 n = 113 | | | | Sh vs. Sh Old ΔAO Inflexible P=0.3167 | | |
| | | Sh Aged ΔAO Inflexib | 0.43 ± 0.15 n = 131 | | | | Sh Old vs. Sh Old ΔAO Flexible ****P=<0.0001 | | |
| | | | | | | | Sh Old vs. Sh Old ΔAO Inflexible P=0.8829 | | |
| | | | | | | | Sh Old ΔAO Flexible vs. Sh Old ΔAO Inflexible ****P=<0.0001 | | |
| | DAGLα LP / Clusters | | | | | | Dunn's multiple comparisons test | | Kruskal-Wallis test |
| | | Sh | 1 ± 0.21 n = 23 | | | p=0,1973 | Sh vs. Sh Old P=>0.9999 | | |
| | | Sh Aged | 0.77 ± 0.09 n = 27 | | | | Sh vs. Sh Old ΔAO Flexible P=>0.9999 | | |
| | | Sh Aged ΔAO Flexibl | 1.21 ± 0.16 n = 59 | | | | Sh vs. Sh Old ΔAO Inflexible P=>0.9999 | | |
| | | Sh Aged ΔAO Inflexib | 0.65 ± 0.11 n = 11 | | | | Sh Old vs. Sh Old ΔAO Flexible P=>0.9999 | | |
| | | | | | | | Sh Old vs. Sh Old ΔAO Inflexible P=>0.9999 | | |
| | | | | | | | Sh Old ΔAO Flexible vs. Sh Old ΔAO Inflexible P=0.5424 | | |
| **Fig. 5F** | Cb1 LP / μm2 | | | | | | Dunn's multiple comparisons test | | Kruskal-Wallis test |
| | | Sh | 1.00 ± 0.12. n = 77 | | | ****p<0.0001 | Sh vs. Sh_Old *P=0.0281 | | |
| | | Sh Aged | 0.58 ± 0.07. n = 73 | | | | Sh vs. Sh_Old ΔAO Flexible ****P=<0.0001 | | |
| | | Sh Aged ΔAO Flexibl | 2.49 ± 0.26. n = 73 | | | | Sh vs. Sh_Old ΔAO Inflexible **P=0.0013 | | |
| | | Sh Aged ΔAO Inflexib | 0.43 ± 0.05. n = 55 | | | | Sh_Old vs. Sh_Old ΔAO Flexible ****P=<0.0001 | | |
| | | | | | | | Sh_Old vs. Sh_Old ΔAO Inflexible P=>0.9999 | | |
| | | | | | | | Sh_Old ΔAO Flexible vs. Sh_Old ΔAO Inflexible ****P=<0.0001 | | |
| | Cb1 Clusters / μm2 | | | | | | Dunn's multiple comparisons test | | Kruskal-Wallis test |
| | | Sh | 1.00 ± 0.21. n = 77 | | | ****p<0.0001 | Sh vs. Sh_Old P=0.1582 | | |
| | | Sh Aged | 0.44 ± 0.10. n = 73 | | | | Sh vs. Sh_Old ΔAO Flexible ****P=<0.0001 | | |
| | | Sh Aged ΔAO Flexibl | 2.52 ± 0.28. n = 73 | | | | Sh vs. Sh_Old ΔAO Inflexible *P=0.0456 | | |
| | | Sh Aged ΔAO Inflexib | 0.34 ± 0.11. n = 55 | | | | Sh_Old vs. Sh_Old ΔAO Flexible ****P=<0.0001 | | |
| | | | | | | | Sh_Old vs. Sh_Old ΔAO Inflexible P=>0.9999 | | |
| | | | | | | | Sh_Old ΔAO Flexible vs. Sh_Old ΔAO Inflexible ****P=<0.0001 | | |
| | Cb1 LP / Clusters | | | | | | Dunn's multiple comparisons test | | Kruskal-Wallis test |
| | | Sh | 1 ± 0.14 n = 34 | | | **p=0,0089 | Sh vs. Sh_Old P=>0.9999 | | |
| | | Sh Aged | 0.98 ± 0.19 n = 20 | | | | Sh vs. Sh_Old ΔAO Flexible P=>0.9999 | | |
| | | Sh Aged ΔAO Flexibl | 1.11 ± 0.1 n = 60 | | | | Sh vs. Sh_Old ΔAO Inflexible P=0.0834 | | |
| | | Sh Aged ΔAO Inflexib | 0.57 ± 0.05 n = 11 | | | | Sh_Old vs. Sh_Old ΔAO Flexible P=0.5869 | | |
| | | | | | | | Sh_Old vs. Sh_Old ΔAO Inflexible P=0.5412 | | |
| | | | | | | | Sh_Old ΔAO Flexible vs. Sh_Old ΔAO Inflexible **P=0.0071 | | |
| **Fig. 5G** | | | | | | | | | |
| **Adaptive Index / DGL LP/μm2** | | | | | *P = 0,0367 | | | | |
| | | | | | r2 = 0,7041 | | | | |
| **Adaptive Index / DGL Clusters/μm2** | | | | | *P = 0,0496 | | | | |

| | | | | | | | | | |
|---|---|---|---|---|---|---|---|---|---|
| | | | | | | r2 = 0,6596 | | | |
| **Adaptive Index / DGL LP / Cluster** | | | | | | P = 0,3320 | | | |
| | | | | | | r2 = 0,2313 | | | |
| **Fig. 5H** | | | | | | | | | |
| Adaptive Index / Cb1 LP/µm2 | | | | | | P = 0,0519 | | | |
| | | | | | | r2 = 0,6525 | | | |
| Adaptive Index / Cb1 Clusters/µm2 | | | | | | P = 0,0753 | | | |
| | | | | | | r2 = 0,5878 | | | |
| Adaptive Index / Cb1 LP / Cluster | | | | | | P =0,0785 | | | |
| | | | | | | r2 = 0,6971 | | | |
| **Fig. S1B** | ME min | | | | | | | | Two-way RM ANOVA |
| | Sh | | | n = 23 | Session x Group | F (24. 592) = 1.289 | P=0.1622 | | |
| | ShΔAO | | | n = 17 | Session | F (8. 592) = 40.31 | ****P<0.0001 | | |
| | Ov | | | n = 22 | Group | F (3. 74) = 0.7964 | P=0.4998 | | |
| | OvΔAO | | | n = 16 | | | | | |
| | INP min | | | | Session x Group | F (24. 592) = 1.955 | **P=0.0045 | | Two-way RM ANOVA |
| | Sh | | | n = 23 | Session | F (8. 592) = 5.186 | ****P<0.0001 | | |
| | ShΔAO | | | n = 17 | Group | F (3. 74) = 0.6303 | P=0.5978 | | |
| | Ov | | | n = 22 | | | | | |
| | OvΔAO | | | n = 16 | | | | | |
| **Fig. S1C** | INP min | | | | Contingency x Group | F (1. 31) = 1.422 | P=0.2421 | Sh vs. Sh_Old ΔAO Inflexible P=0.4932 | MD ANOVA |
| | ShΔAO | Positive AO | 0.41 ± 0.16. n = 17 | | Contingency | F (1. 31) = 0.1839 | P=0.6710 | | |
| | | Negative AO | 0.29 ± 0.05. n = 17 | | Group | F (1. 31) = 0.09785 | P=0.7565 | | |
| | OvΔAO | Positive AO | 0.37 ± 0.08. n = 16 | | | | | Ov_ Positive AO vs. Ov_ Negative AO P=0.7925 | |
| | | Negative AO | 0.44 ± 0.09. n = 16 | | | | | | |
| **Fig. S1D** | Sh | | | 80,83 ± 5,824 n = 9 | Group | F (3, 35) = 5,207 | P = 0,6530 | | One-way ANOVA |
| | ShΔAO | | | 78,61 ± 2,811 n = 9 | | | | | |
| | Ov | | | 87,02 ± 6,968 n = 9 | | | | | |
| | OvΔAO | | | 84,68 ± 3,800 n = 12 | | | | | |
| **Fig. S1E** | ANPmin | | | | | | | | |
| | Sh | | | n = 12 | Session x Group | F (24. 392) = 1.083 | P=0.3605 | | Two-way RM ANOVA |
| | ShΔAO | | | n = 16 | Session | F (8. 392) = 83.22 | ****P<0.0001 | | |
| | Ov | | | n = 14 | Group | F (3. 49) = 0.5418 | P=0.6559 | | |
| | OvΔAO | | | n = 11 | | | | | |
| **Fig. S1F** | ANPmin | | | | | | | | |
| | ShΔAO | 5 min | | 0.96 ± 0.09. n = 16 | Time x Group | F (5. 125) = 1.810 | P=0.1155 | | Two-way ANOVA |
| | | 10 min | | 0.77 ± 0.08. n = 16 | Time | F (5. 125) = 8.992 | ****P<0.0001 | | |
| | | 15 min | | 0.61 ± 0.08. n = 16 | Group | F (1. 25) = 29.84 | ****P<0.0001 | | |
| | | 20 min | | 0.49 ± 0.08. n = 16 | | | | | |
| | | 25 min | | 0.44 ± 0.07. n = 16 | | | | | |
| | | 30 min | | 0.36 ± 0.07. n = 16 | | | | | |
| | OvΔAO | 5 min | | 1.16 ± 0.10. n = 11 | | | | | |
| | | 10 min | | 1.18 ± 0.09. n = 11 | | | | | |
| | | 15 min | | 1.15 ± 0.08. n = 11 | | | | | |
| | | 20 min | | 1.09 ± 0.07. n = 11 | | | | | |
| | | 25 min | | 1.04 ± 0.18. n = 11 | | | | | |
| | | 30 min | | 0.84 ± 0.11. n = 11 | | | | | |
| | ANPmin | | | | | | | | |
| | ShΔAO | Positive AO | | 18.71 ± 1.91. n = 16 | Group x A-O Contingency | F (1. 25) = 15.50 | ***P=0.0006 | Sh_Positibve AO vs. Sh_Negative AO ****P<0.0001 | MD Anova |
| | | Negative AO | | 11.75 ± 1.69. n = 16 | Group | F (1. 25) = 2.784 | P=0.1077 | | |
| | | | | | A-O Contingency | F (1. 25) = 12.16 | **P=0.0018 | Ov_ Positive AO vs. Ov_ Negative AO P=0.9483 | |
| | OvΔAO | Positive AO | | 19.66 ± 2.57. n = 11 | | | | | |
| | | Negative AO | | 20.08 ± 2.16. n = 11 | | | | | |
| **Fig. S1G** | Obtained reinforcers | | | | | | | | |
| | ShΔAO | Positive AO | | 0.38 ± 0.01. n = 16 | A-O Contingency x Group | F (1. 25) = 6.071 | *P=0.0210 | Sh_ Positive AO vs. Ov_ Positive AO P=0.9736 | |
| | OvΔAO | Positive AO | | 0.39 ± 0.01. n = 11 | A-O Contingency | F (1. 25) = 24.38 | ****P<0.0001 | | |
| | | | | | Group | F (1. 25) = 6.248 | *P=0.0194 | Sh_ Negative AO vs. Ov_ Negative AO **P=0.0020 | |
| | ShΔAO | Negative AO | | 0.29 ± 0.05. n = 16 | | | | | |
| | OvΔAO | Negative AO | | 0.11 ± 0.04. n = 11 | | | | | |
| **Fig. S1H** | ME min | | | | | | | | Two-way RM ANOVA |
| | Sh | | | n = 12 | Session x Group | F (24. 392) = 1.121 | P=0.3167 | | |
| | ShΔAO | | | n = 16 | Session | F (8. 392) = 6.569 | ****P<0.0001 | | |
| | Ov | | | n = 14 | Group | F (3. 49) = 0.7830 | P=0.5092 | | |
| | OvΔAO | | | n = 11 | | | | | |
| | INP min | | | | Session x Group | F (24. 392) = 1.680 | *P=0.0246 | | Two-way RM ANOVA |

| | | | | | | | | | | |
|---|---|---|---|---|---|---|---|---|---|---|
| | Sh | | | n = 12 | Session | F (8. 392) = 2.308 | *P=0.0199 | | | |
| | ShΔAO | | | n = 16 | Group | F (3. 49) = 0.5313 | P=0.6630 | | | |
| | Ov | | | n = 14 | | | | | | |
| | OvΔAO | | | n = 11 | | | | | | |
| **Fig. S1I** | INP min | | | | Group x A-O Contingency | F (1. 25) = 4.424 | P=0.0457 | Sh vs. Sh_Old ΔAO Inflexible P=0.9451 | | Wilcoxon Paired Test |
| | ShΔAO | Positive AO | 0.18 ± 0.04. n = 16 | | Group | F (1. 25) = 1.472 | P=0.2364 | | | |
| | | Negative AO | 0.17 ± 0.03. n = 16 | | A-O Contingency | F (1. 25) = 2.954 | P=0.0980 | | | |
| | OvΔAO | Positive AO | 0.22 ± 0.08. n = 11 | | | | | Ov_ Positive AO vs. Ov_ Negative AO *P=0.0398 | | |
| | | Negative AO | 0.34 ± 0.11. n = 11 | | | | | | | |
| **Fig. S2B** | ME min | | | | | | | | | Two-way RM ANOVA |
| | Sh | | | n = 10 | Session x Group | F (16, 272) = 0.4597 | P=0.9638 | | | |
| | Sh_Aged | | | n = 10 | Session | F (8, 272) = 7.967 | ****P<0.0001 | | | |
| | Sh_Aged ΔAO | | | n = 17 | Group | F (2, 34) = 0.6414 | P=0.5328 | | | |
| | INP min | | | | | | | | | Two-way RM ANOVA |
| | Sh | | | n = 10 | Session x Group | F (16, 272) = 1.377 | P=0.1525 | | | |
| | Sh_Aged | | | n = 10 | Session | F (8, 272) = 4.535 | ****P<0.0001 | | | |
| | Sh_Aged ΔAO | | | n = 17 | Group | F (2, 34) = 0.4036 | P=0.6711 | | | |
| **Fig. S2C** | INP min | | | | | | | | | MD ANOVA |
| | Sh_Aged ΔAO Flex | Positive AO | 0.14 ± 0.03. n = 10 | | Group x A-O Contingency | F (1. 15) = 5.118 | *P=0.0390 | Control - Reversal | | |
| | | Negative AO | 0.11 ± 0.02. n = 7 | | Group | F (1. 15) = 3.145 | P=0.0964 | Short GD | P=0.9985 | |
| | | | | | A-O Contingency | F (1. 15) = 4.837 | *P=0.0440 | Short Hab | *P=0.0215 | |
| | Sh_Aged ΔAO Infle | Positive AO | 0.14 ± 0.05. n = 10 | | | | | | | |
| | | Negative AO | 0.58 ± 0.25. n = 7 | | | | | | | |
| | Obtained reinforcers | | | | | | | | | MD ANOVA |
| | Sh_Aged ΔAO Flex | Positive AO | 0.35 ± 0.01. n = 10 | | A-O Contingency x Group | F (1, 15) = 49,52 | ****P<0,0001 | Short GD - Short Hab | | |
| | Sh_Aged ΔAO Infle | Positive AO | 0.39 ± 0.01. n = 7 | | A-O Contingency | F (1, 15) = 0,6640 | P=0,4279 | Control | p=0,772 | |
| | | | | | Group | F (1, 15) = 47,33 | ****P<0,0001 | Reversal | ****p<0,0001 | |
| | Sh_Aged ΔAO Flex | Negative AO | 0.60 ± 0.06. n = 10 | | | | | | | |
| | Sh_Aged ΔAO Infle | Negative AO | 0.07 ± 0.02. n = 7 | | | | | | | |
